# Pumping Iron: A Multi-omics Analysis of Two Extremophilic Algae Reveals Iron Economy Management

**DOI:** 10.1101/2023.02.09.527888

**Authors:** Lital Davidi, Sean D. Gallaher, Eyal Ben-David, Samuel O. Purvine, Thomas L. Filmore, Carrie D. Nicora, Rory J. Craig, Stefan Schmollinger, Sanja Roje, Crysten E. Blaby-Haas, Robert P. Auber, Jennifer Wisecaver, Sabeeha S. Merchant

## Abstract

Marine algae are responsible for half of the world’s primary productivity, but this critical carbon sink is often constrained by insufficient iron. One species of marine algae, *Dunaliella tertiolecta*, is remarkable for its ability to maintain photosynthesis and thrive in low-iron environments. A related species, *Dunaliella salina* Bardawil, shares this attribute but is an extremophile found in hyper-saline environments. To elucidate how algae manage their iron requirements, we produced high-quality genome assemblies and transcriptomes for both species to serve as a foundation for a comparative multi-omics analysis. We identified a host of iron-uptake proteins in both species, including a massive expansion of transferrins and a novel family of siderophore-iron uptake proteins. Complementing these multiple iron-uptake routes, ferredoxin functions as a large iron reservoir that can be released by induction of flavodoxin. Proteomic analysis revealed reduced investment in the photosynthetic apparatus coupled with remodeling of antenna proteins by dramatic iron-deficiency induction of TIDI1, an LHCA-related protein found also in other chlorophytes. These combinatorial iron scavenging and sparing strategies make *Dunaliella* unique among photosynthetic organisms.

**Significance Statement:** Despite their small size, microalgae play a huge role in CO_2_ uptake via photosynthesis, and represent an important target for climate crisis mitigation efforts. Most photosynthesis proteins require iron as a co-factor so that insufficient iron often limits algal CO_2_ sequestration. With this in mind, we examined a genus of microalgae called *Dunaliella* that is exceptionally well-adapted to low iron environments. We produced complete genomes, transcriptomes, and proteomes for two species of *Dunaliella* that hail from radically different environments: one from coastal ocean waters and the other from salt flats. We identified dozens of genes and multiple, complementary strategies that both species utilize for iron-uptake and management that explain *Dunaliella’s* remarkable ability to thrive on low iron.

## Introduction

The atmospheric release of CO_2_ due to human activity represents a grave threat to world ecosystems and human well-being. An important counter-balance to rising greenhouse gas concentrations is primary productivity, wherein CO_2_ is taken up by photosynthetic organisms and converted to organic compounds. Approximately one half of primary productivity on Earth is due to the photosynthetic activity of marine algae (1). Unfortunately, this critical carbon sink is constrained by insufficient Fe in approximately one third of the Earth’s oceans (2). As evidenced by mesoscale Fe-addition experiments, seeding the water in these high nutrient-low chlorophyll regions with Fe is sufficient to induce blooms of phytoplankton (3). Thus, a greater understanding of the role of Fe uptake and homeostasis in algae represents an important research aim for mitigation of the climate crisis.

Not all algae are equally adept at acquiring Fe from their environment. While Fe is the fourth most abundant element in the Earth’s crust, it has poor bioavailability (4, 5). In our oxygenated world, Fe reacts with oxygen, to form insoluble oxyhydroxides that become inaccessible to many aquatic organisms or energetically costly to access. One genus of green algae that is exceptional at Fe uptake is *Dunaliella*. This genus comprises motile, unicellular green algae of the chlorophyte lineage, which diverged ∼500 Mya (6) from the lineage that includes the reference chlorophycean alga, *Chlamydomonas reinhardtii. Dunaliella* species are extremely halotolerant, as well as tolerant of extremes of high light, temperature and a wide range of pH (7). *Dunaliella* species are cosmopolitan in their distribution, including coastal sea waters, salty lakes, and even salt flats (Fig. 1A). *Dunaliella* species lack a rigid cell wall. Instead, they achieve osmotic balance with their saline environment by synthesizing and storing high concentrations of glycerol in their cytoplasm as an osmolyte (8).

**Fig. 1.**
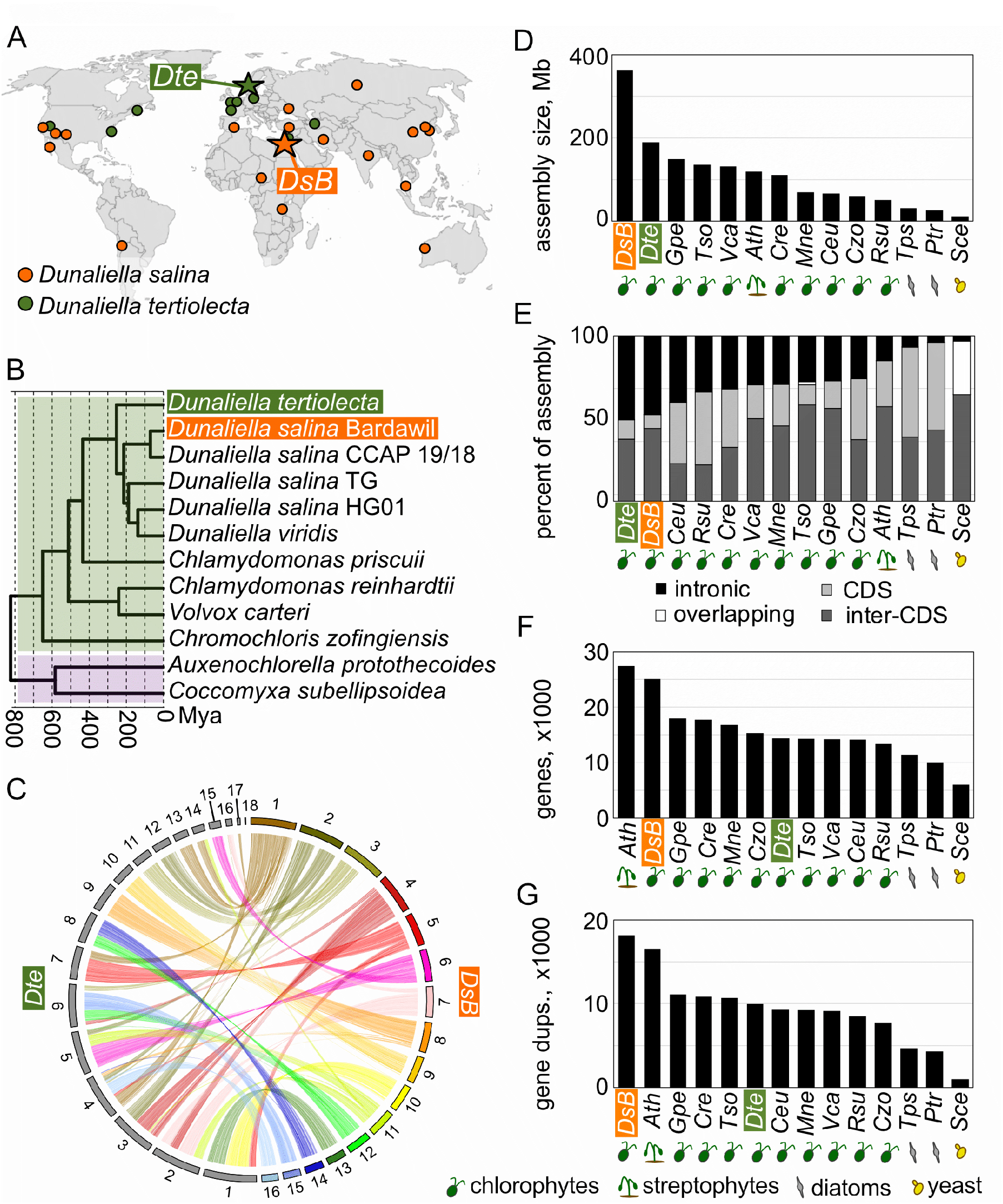
*D. salina* Bardawil and *D. tertiolecta* have large genomes due to large introns and widespread gene copy number expansion. (**A**) The geographic origins of many field isolates of *D. salina* (orange) and *D. tertiolecta* (green) are shown. The strains used in this work, *D. tertiolecta* UTEX LB 999 and *D. salina* Bardawil UTEX LB 2538, are indicated by stars. (**B**) A chronogram shows the divergence times between several isolates of *Dunaliella* and other green algae from class Chlorophyceae (green) and Trebouxiophyceae (lavender). The chronogram is based on the concatenated polypeptide sequences of 104 universal single copy orthologs (USCOs) shared among all 12 species (see Methods). (**C**) A circos plot shows the synteny between the 16 chromosomes of *D. salina* Bardawil (right half) and the 18 scaffolds of *D. tertiolecta* (left half), and demonstrates the 1:1 relationship between the two assemblies. (**D**) The size of each genome assembly is presented as a bar plot arranged from largest to smallest. A three-letter code identifies each species as follows: *DsB* = *D. salina* Bardawil, *Dte* = *D. tertiolecta, Gpe* = *Gonium pectorale, Tso* = *Tetrabaena socialis, Vca* = *Volvox carteri, Ath* = *Arabidopsis thaliana, Cre* = *Chl. reinhardtii, Mne* = *Monoraphidium neglectum, Ceu* = *Chlamydomonas eustigma, Czo* = *Chromochloris zofingiensis, Rsu* = *Raphidocelis subcapitata, Tps* = *Thalassiosira pseudonana, Ptr* = *Phaeodactylum triconutum*, and *Sce* = *Saccharomyces cerevisiae*. Species identifiers are decorated with a symbol indicating their phylogenetic lineage according to the table at the bottom-right of the figure. (**E**) For the same species, the percentage of each genome assembly that is protein coding sequence (CDS), intronic, inter-CDS sequence, or some overlap of CDS and/or intronic, is indicated by the color-coded fractions of each bar. Species are arranged from the most to the least intronic. Further analysis of the contributions of introns is detailed in SI Appendix, Fig. S3. (**F**) For the same species, the total number of annotated protein coding genes is presented as a bar plot arranged from most to fewest genes. (**G**) Gene duplications for the lineages leading to each species were computationally predicted. The number of gene duplications is plotted as a bar plot arranged in decreasing order.

*Dunaliella* species have been explored in a number of biotechnological applications due to their ability to thrive in harsh environments that suppress the growth of other species (9). Using only the widely available inputs of sunlight, seawater and CO_2,_ *Dunaliella* is capable of robust growth and can be harnessed to produce various, useful products without competing for valuable resources (e.g. fertile soil and freshwater) necessary for other more traditional forms of agriculture. These features make *Dunaliella* an appealing platform for the sustainable production of many important bioproducts, including glycerol and carotenoids (10, 11).

In this work, we focused on two species of *Dunaliella* that live in radically different environments. The first, *D. tertiolecta*, is commonly found in coastal sea waters worldwide (Fig. 1A). It is under development for the production of biofuels because it is capable of accumulating large quantities of neutral lipids that can be refined into biodiesel (12). The second species, *D. salina*, is found in hypersaline environments and salt flats. *D. salina* has many of the same characteristics as *D. tertiolecta*, making it an attractive platform for producing bioproducts. For example, the strain of *D. salina* used in this work is the richest natural source of β-carotene, a highly valuable commercial product (9, 13). Having been isolated from a salt pond near the Bardawil Lagoon, this strain is sometimes called *D. bardawil*, and will be referred to as *D. salina* Bardawil throughout this work.

Much like their ability to tolerate extremes of temperature and salt, *Dunaliella* species are exceptional at maintaining Fe homeostasis in environments where the bioavailable Fe is growth limiting to other species. Tolerance to high salt and low Fe bioavailability may have co-evolved since extreme saline conditions drastically reduce Fe solubility. Remarkably, these algae are able to maintain photosynthetic capacity under polyextreme conditions without a major impact on growth (14), suggesting the existence of unique functionality not found in other photosynthetic organisms. Here we have employed a multi-omics strategy to elucidate how *D. tertiolecta* and *D. salina* Bardawil manage environmental extremes, especially low Fe. As a foundation for this work, we generated high-quality, chromosome-scale genome assemblies and transcriptomes for both species. With transcriptomics and proteomics analyses, we identified the key proteins that these species rely on for their exceptional ability to thrive in low Fe and other harsh conditions. We elucidated the mechanisms that *Dunaliella* deploys for maintaining growth in low Fe and related these to strategies used by other plants and by diatoms. The annotated genomes coupled with transcriptomic and proteomic data that accompany this work will provide a solid foundation to the community for continuing discoveries on extremophilic algae and beyond.

## Results

### Genomics, transcriptomics, and proteomics of *D. tertiolecta* and *D. salina* Bardawil reveal keys genes important for Fe homeostasis

To elucidate how these *Dunaliella* species evolved to manage their Fe quota (SI Appendix, Fig. S1), we chose a systems biology approach; a strategy best employed when a high-quality genome assembly and a complete set of gene annotations can provide a solid foundation for the analysis. Using a combination of complementary techniques, we generated chromosome-scale assemblies for the haploid genomes of both *D. salina* Bardawil and *D. tertiolecta* (Fig. 1). We found that these two extremophilic *Dunaliella* species, which we estimate diverged ∼253 Mya (Fig. 1B), have exceptionally large genomes relative to other mesophilic chlorophycean algae (Fig. 1D and SI Appendix, text). For the *Dunaliella* genome assemblies presented here, we attribute their considerable size to a combination of being rich in repetitive sequences and transposable elements (SI Appendix, text, Fig. S2 and Table S1), having very large introns (Fig. 1E and SI Appendix, Fig. S3), and to widespread gene copy number expansion (Fig. 1F-G). This last point proved to be an important feature of Fe homeostasis in both *Dunaliella* species, as we note below.

Next, we performed transcriptomics and proteomics on both species grown in defined media with and without added Fe to identify the key Fe-homeostasis genes (Fig. 2). While the vast majority of *D. salina* Bardawil’s ∼25,000 genes were minimally affected by the change in available Fe, the expression of a few hundred genes was radically altered, increasing by as much as 10^3^-fold in the –Fe samples. In *D. salina* Bardawil, we identified 80 up-regulated and 50 down-regulated (Fig. 2A) differentially expressed genes (DEGs). In *D. tertiolecta*, which was more significantly impacted by –Fe growth, we identified 169 up-regulated and 239 down-regulated DEGs (Fig. 2B). Most of these Fe-regulated genes had orthologs in both species, which suggests *Dunaliella*’s Fe homeostasis mechanisms predate the split of these two species ∼253 Mya. Included in the group of genes that are highly up regulated in –Fe are all the components of a reductive Fe-uptake pathway, such as a Ferroxidase (FOX1) and a Ferric reductase (FRE1A) (SI Appendix, Fig. S4). The down-regulated genes included components of the photosynthetic apparatus, and the Fe-storage protein, Ferritin (FER1). In the following sections, we present some of the more unexpected discoveries.

**Fig. 2.**
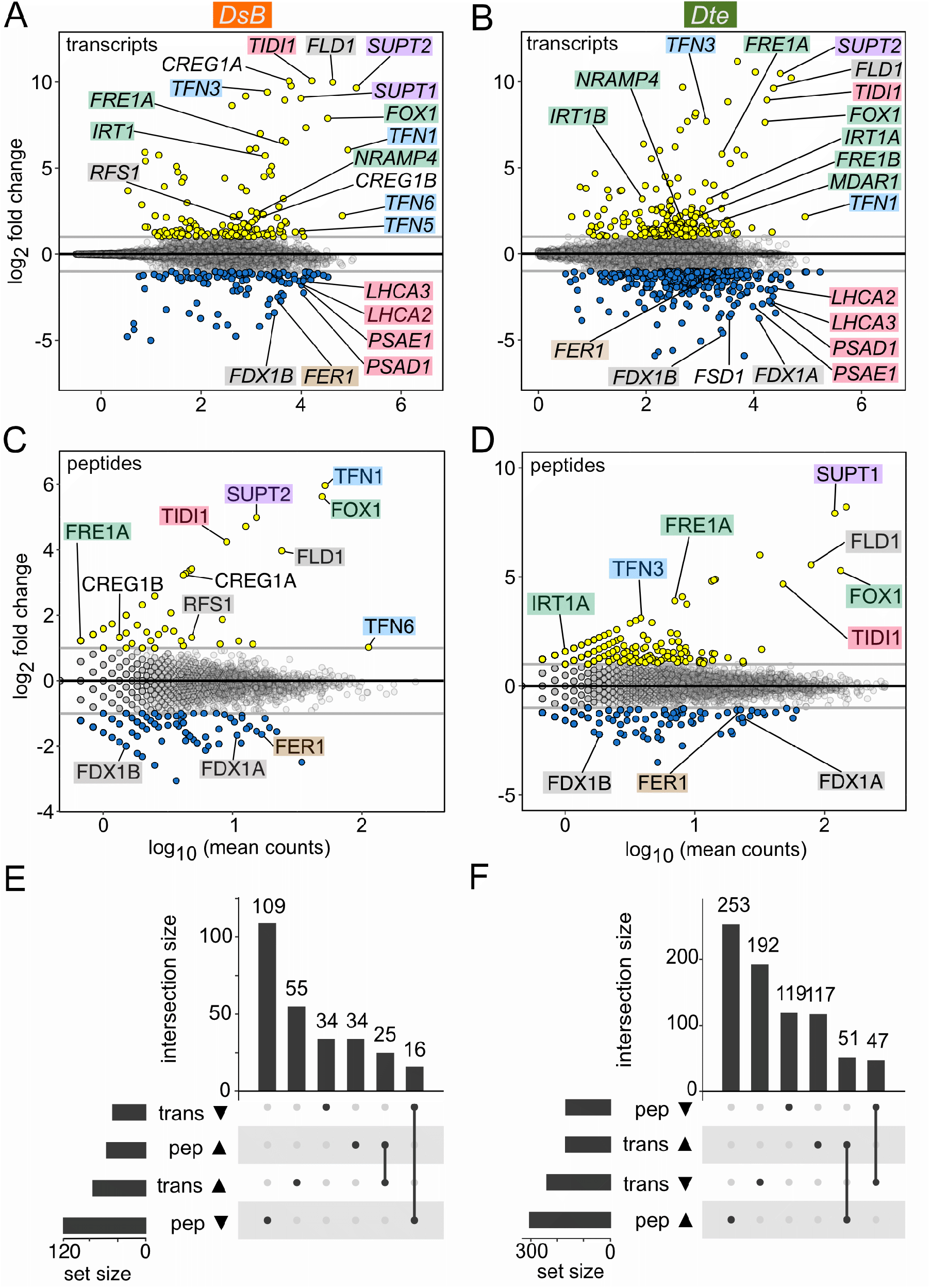
Dynamic gene expression in *D. salina* Bardawil and *D. tertiolecta* in response to –Fe. Cultures of *D. salina* Bardawil and *D. tertiolecta* were grown in media +/–Fe and gene expression was analyzed via transcriptomics and proteomics. (**A**) An MA plot demonstrates differentially expressed genes (DEGs) for *D. salina* Bardawil in –Fe relative to +Fe: up regulated in yellow and down regulated in blue. A log_2_ fold-change=1 threshold is indicated by horizontal gray lines. Select DEGs are labeled in the figure with the following color scheme: green = reductive Fe uptake proteins, blue = transferrins, lavender = siderophore uptake proteins, pink = photosystem components, brown = ferritin, and gray = Fe-sparing related. (**B**) The same analysis presented in panel A was performed here on *D. tertiolecta*. (**C**) MA plot of differentially detected spectral counts for proteins from *D. salina* Bardawil cultures grown without Fe versus with Fe. Proteins were considered up (yellow) or down (blue) if the log_2_ fold change of spectral counts plus a pseudo count of 1 was >1. Select differentially detected proteins are labeled using the same color scheme as in panel A and B. (**D**) The same analysis was performed here on *D. tertiolecta*. (**E**) An upset plot demonstrates the overlap between differentially detected transcripts and differentially detected proteins for *D. salina* Bardawil in response to –Fe. (**F**) The same analysis was performed here for *D. tertiolecta*.

### There has been a major expansion of Fe-binding Transferrin family proteins in *Dunaliella*

When we examined the most highly Fe-regulated genes, we found a large number of previously unknown Transferrin-family (Tf) proteins in both *Dunaliella* species. Tf proteins were first identified in mammals as soluble glycoproteins with two Fe^+3^ ion-binding domains that mediate Fe transport into cells by binding a cell surface receptor and subsequent endocytosis (15). While two Tf-family proteins had been identified in *D. tertiolecta* previously (16, 17), we discovered a much larger expansion of Tf-encoding genes in the *Dunaliella* lineage (SI Appendix, Fig. S5), resulting in seven Tf-encoding genes in *D. tertiolecta* and six in *D. salina* Bardawil (Fig. 3A). Estimation of the divergence times of the different *Dunaliella* Tf proteins (SI Appendix, Fig. S5) indicates that Tf gene expansion in the *Dunaliella* lineage occurred at multiple points after the Neoproterozoic oxygenation event. This period was marked by increased levels of O_2_ leading to oxidation of environmental Fe; factors that would place a premium on ferric iron-uptake mechanisms.

**Fig. 3.**
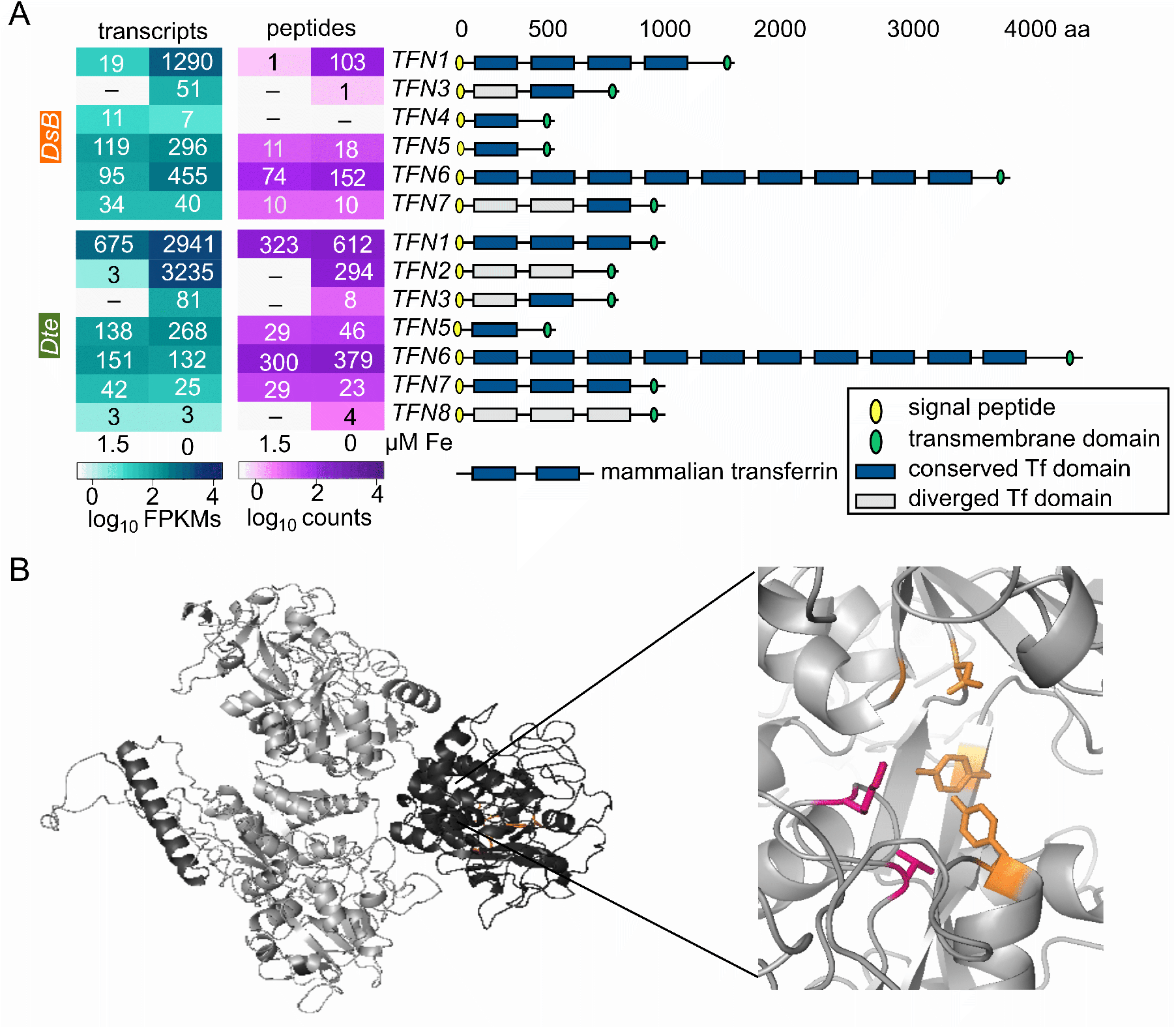
Expansion of Transferrin-encoding genes in *Dunaliella* help manage Fe scarcity. (**A**) On the left, a heatmap of transcript abundance estimates as log_10_-transformed FPKMs and protein abundance estimates as log_10_-transformed spectral counts is presented for the six transferrins of *D. salina* Bardawil, and the seven transferrins of *D. tertiolecta* from cultures grown +/– Fe, as indicated. To the right, a cartoon demonstrates significant features of each predicted protein as described by the legend. A Tf domain was considered conserved (dark blue) if it retains five key residues involved in Fe and anion binding, or diverged (gray) if not. A protein model of a canonical two-domain, soluble human Tf is included for reference. (**B**) Structure prediction of *D. tertiolecta* TFN1 shows a three-domain protein. An inset panel magnifies residues that are expected to be involved in Fe binding (orange) and anion binding (pink). See SI Appendix, Table S5 for details on conservation of Fe and anion binding residues.

The classically defined Tf proteins in other species predominantly have one or two Tf domains. In contrast, we found that the Tf proteins expressed in *Dunaliella* were highly variable in size, with as few as one and as many as ten Tf motifs (Fig. 3A). All 13 *Dunaliella* proteins were predicted to have a trans-membrane domain, as well as a signal peptide and one or more glycosylation sites, supporting a role for the Tf proteins on the cell surface with their Fe-binding domains extending outside the cell. Thus, it is likely that the *Dunaliella* Tf proteins have the dual function of Fe binding and internalization, which contrasts with the soluble Tf proteins of metazoans that must bind a receptor for cellular Fe uptake. Despite these differences relative to the classically defined Tfs of metazoans, the conservation of Fe and anion binding residues (SI Appendix, Table S2) and the predicted structures of the *Dunaliella* Tf domains validates that these are true Tf proteins (SI Appendix, Fig. S6).

Transcriptomic and proteomic analyses of the *Dunaliella* Tfs shows very high levels of expression and protein abundance for many of the Tfs. The most abundant ones are dramatically up regulated in the low Fe condition (Fig. 3A), suggesting an important role for these Tf proteins in Fe uptake. In a survey of all 44 *Dunaliella* Tf domains (from 13 Tf proteins), we observed that domains generally either retain all Fe-binding amino acid residues (18) (shown as “conserved” dark blue boxes in Fig. 3A) or retain few of the conserved residues (shown as “diverged” gray boxes). The Tf proteins that have retained most or all of the Fe-binding residues were also the most highly expressed in response to low Fe. Combined with the lack of Fe-responsive expression, the divergent Tf proteins, such as *D. tertiolecta* TFN8, may be becoming pseudogenes or have developed a different, novel function.

### A family of proteins of unknown function may mediate Siderophore-Fe uptake in *Dunaliella*

In a previous analysis of *D. tertiolecta* (incorrectly reported in that work as *D. salina*), an unknown plasma membrane-bound glycoprotein was isolated from Fe-limited cells (19). This protein was only detected under Fe-limitation conditions and was physically associated with FOX1, TFN1, and TFN2. At that time, the authors were unable to find similarity between this glycoprotein and any known proteins and named it “p130B” based on its 130 kDa molecular weight. In the present survey, we identified the *D. tertiolecta* p130B gene as one of a family of genes with multiple homologs in both *Dunaliella* species: two in *D. salina* Bardawil and three in *D. tertiolecta*.

Transcription of all five of these *Dunaliella* p130B-like genes, which we have named *SUPT1* - *SUPT4* (see below), was strongly up regulated in response to Fe limitation (Fig. 4A). Protein abundances were similarly increased in the –Fe condition for all except *D. salina* Bardawil SUPT1; an observation with we attribute to a recent nonsense mutation in the corresponding gene (SI Appendix, text and Fig. S7). Similar to the Tfs, all five of the *SUPT* genes were predicted to encode a signal peptide, a transmembrane helix at the C-terminus, and one or more glycosylation sites (Fig. 4A), which is consistent with a role for these proteins on the cell surface. Structure prediction of SUPT1 from *D. tertiolecta* revealed two adjacent seven-bladed β-propeller domains and a transmembrane domain, which is consistent with a cell surface-located, ligand-binding protein (Fig. 4C).

**Fig. 4.**
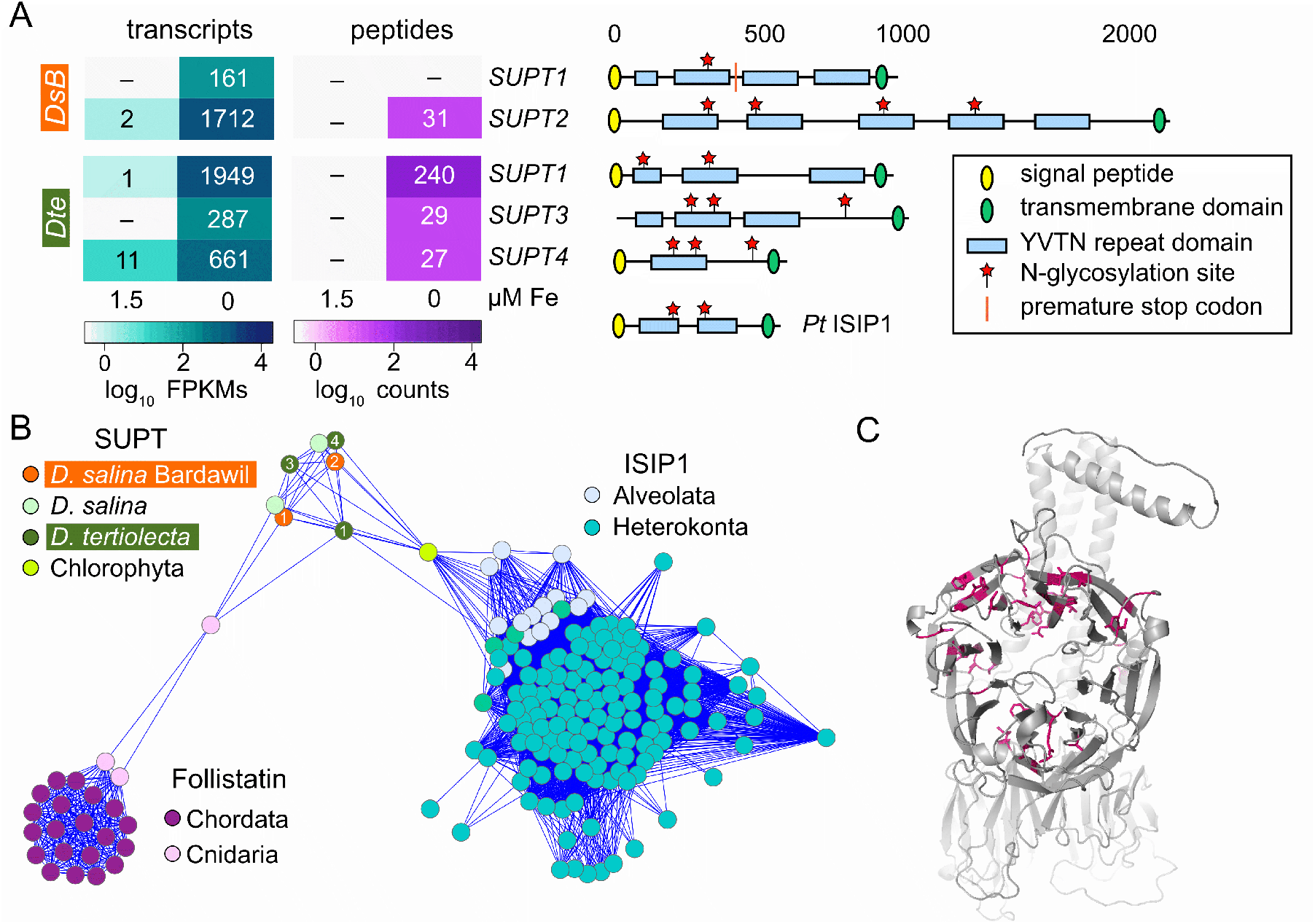
P130B-family proteins may facilitate siderophore-Fe uptake in *Dunaliella* species. (**A**) Heatmaps for transcript and protein levels of siderophore uptake protein 1 (*SUPT1)* - *SUPT4* in *D. salina* Bardawil and *D. tertiolecta* grown in media +/– Fe. To the right of each *SUPT* gene symbol is a cartoon of the predicted protein structure. Predicted features are presented as described in the accompanying legend. A single stop codon in *D. salina* Bardawil *SUPT1* breaks up an otherwise conserved open reading frame, and is indicated here by a red line (SI Appendix). (**B**) A protein similarity network of *Dunaliella* SUPT proteins and likely chlorophyte orthologs, ISIP1 proteins from diatoms, and metazoan follistatins. The similarity network was generated using blastp comparison of all-vs-all on over 100 sequences using E-value of 1 × 10^−25^. (**C**) The structure of *D. tertiolecta* SUPT1 was predicted to consist of two 7-blade propeller domains, one facing the viewer and the second perpendicular to it, and an alpha helix transmembrane domain. The 50 aa that are conserved amongst all *Dunaliella* SUPT proteins and their *Chlamydomonas schloesseri* orthologs are marked in pink.

To identify the role of SUPT-family proteins in coping with Fe-deficiency stress, we looked for similar proteins with known function. The *Dunaliella* SUPT proteins share similarity with metazoan Follistatin, which is a cell surface-bound, ligand-binding glycoprotein. More importantly, the SUPTs share similarity with iron starvation-induced protein 1 (ISIP1) in diatoms such as *Phaeodactylum tricornutum* (Fig. 4B). ISIP1 mediates siderophore-Fe uptake via endocytosis (20). Like *D. tertiolecta* SUPT1, we predict that ISIP1 in *P. tricornutum* has a seven-bladed β-propeller structure. The similarities in sequence and predicted structure hint that ISIP1 and SUPT1 may share a function in siderophore-Fe uptake. In support of this hypothesis, *D. salina* Bardawil can utilize siderophore-bound Fe from both microbial siderophores and synthetic chelates (21), but a mechanism for this uptake was not identified until this work.

In summary, all five *Dunaliella SUPT* genes are up regulated in response to low Fe, encode cell surface proteins with a structure commonly found in ligand-binding proteins, and have sequence and structural similarity to siderophore-Fe importing proteins in diatoms. From these observations, we propose that the SUPT family of proteins functions as the previously unknown mediator of siderophore-bound Fe uptake in *Dunaliella*. Based on this function, we have assigned the name Siderophore uptake protein (SUPT) to this family of genes.

### Quantitative proteomics elucidates the Fe-sparing benefit of replacing ferredoxin with flavodoxin in *Dunaliella*

While increased assimilation of an essential mineral nutrient, such as Fe or Cu, is a first line of defense in the face of deficiency, elemental sparing is another (22–24). During sparing, the cell replaces a protein that is dependent on the missing element with an isofunctional protein that does not (25). Typically, this mechanism operates on abundant proteins leading to significant reduction in the demand for the missing element. The replacement of FeS-containing ferredoxin, which functions in various anaerobic bioenergetic pathways, by flavin-containing flavodoxin under poor iron nutrition was noted decades ago in *Clostridium spp*. (26, 27). This switch also occurs in the photosynthetic apparatus of cyanobacteria (28, 29), where the dominance of either protein is a classic biomarker for the iron status of the ocean (30). The operation of this ferredoxin/flavodoxin switch is not well-investigated in eukaryotic algae. In fact, the flavodoxin gene is absent in plants and in *Chlamydomonas* species (31) leading to the dogma that this sparing mechanism was not retained after evolution of the chloroplast from the endosymbiont. Nevertheless, analysis of the *Dunaliella* genomes revealed *FLD1*, which encodes a type 2 flavodoxin (32), and a pair of paralogous ferredoxin-encoding genes (*FDX1A* and *FDX1B*) in both *D. tertiolecta* and *D. salina* Bardawil. The *FDX1* genes are highly expressed in +Fe, with *FDX1A* being dominant, and dramatically down regulated in –Fe (Fig. 5A), while expression of *FLD1* was counter-regulated. Curiously, down regulation of *FDX1A* occurred at the level of transcription in *D. tertiolecta* but at the level of protein abundance in *D. salina* Bardawil (Fig. 5A), speaking perhaps to independent evolution of the regulatory circuit that controls sparing.

**Fig. 5.**
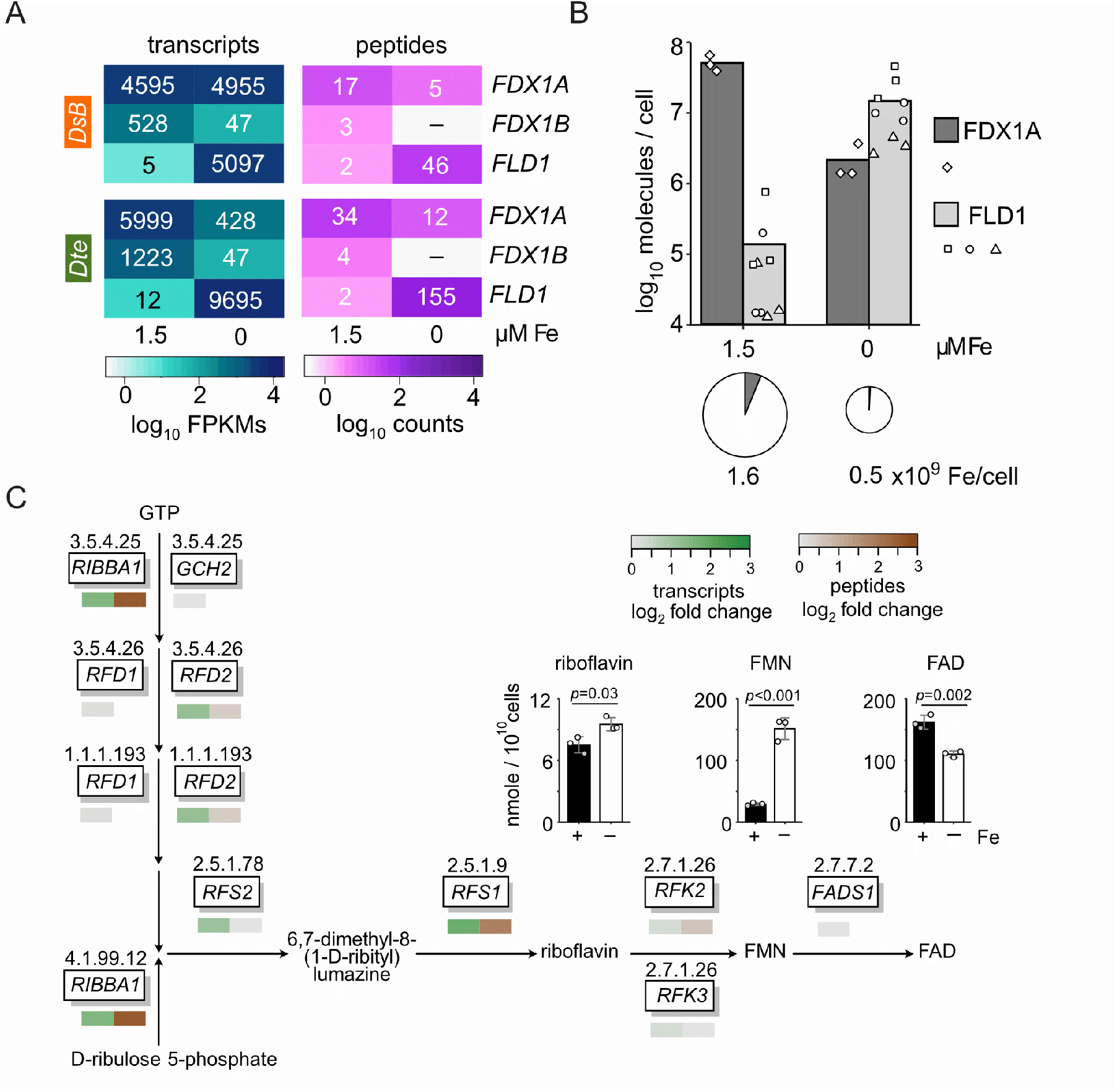
*Dunaliella* replaces ferredoxin with flavodoxin as an Fe-sparing mechanism. (**A**) Transcript and protein abundances for *FDX1A, FDX1B* and *FLD1* are presented for *D. tertiolecta* and *D. salina* Bardawil cultures grown in media +/– Fe. (**B**) Selected reaction monitoring (SRM) proteomics was used to quantify the number of FDX1A (dark gray) and FLD1 (light gray) molecules in *D. salina* Bardawil cells grown in media +/– Fe. The resulting counts per cell were log_10_ transformed, and plotted as the mean number per cell (top). Triplicate determinations of one peptide for FDX1A (*n* = 3) and three peptides for FLD1 (*n* = 9) are overlaid as white symbols. Below, the total cellular Fe content was quantified by ICP-MS and is indicated by the relative size of each pie. The proportion of Fe atoms expected to be bound to FDX1A was calculated assuming two Fe atoms per molecule of FDX1A, and is shown by the dark gray pie slice. (**C**) Components of the flavin biosynthesis pathway are presented along with their products. Heat maps showing the fold change in transcripts (green) and proteins (brown) in –Fe relative to +Fe are shown under each gene. The quantity of three key metabolites, riboflavin, FMN and FAD, were measured from cultures grown +/– Fe (*n* = 3).

Next, we quantified the benefit of replacing FDX with FLD as an Fe-sparing mechanism for eukaryotic organisms. Replacement of FDX1A with FLD1 allows the cell to free up two Fe atoms per molecule of FDX1A. To evaluate the impact of replacement on a per cell basis, we quantified the two proteins in *D. salina* Bardawil by selected reaction monitoring (SRM) proteomics (33). FDX1A, at an abundance of 5.1 × 10^7^ molecules / cell, decreased by 23-fold in the low Fe condition to 2.2 × 10^6^ molecules / cell, while FLD1 increased 106-fold to an abundance of 1.5 × 10^7^ molecules / cell (Fig. 5B). Thus, a *D. salina* Bardawil cell can free up 1 × 10^8^ atoms of Fe solely by replacing FDX1A with FLD1. Concurrently, the Fe quota of the algal cell decreased three-fold from 1.5 to 0.49 × 10^9^ atoms per cell in +Fe versus –Fe, which translates to an Fe allocation of 6% to FDX1A in the Fe-replete situation versus only 0.9% in low Fe.

The metabolic cost of this switch (besides perhaps reduced electron transfer rates by the substitute protein (34)) is a substantial draw on the flavin cofactor pool (35). Among the DEGs, we noted increased expression of key enzymes in the flavin biosynthetic pathway of *D. salina* Bardawil (Fig. 5C), including the dual-function riboflavin biosynthesis protein (RIBBA1), which initiates the flavin biosynthesis pathway and riboflavin synthase 1 (RFS1), which converts 6,7-dimethyl-8- (1-D-ribityl) lumazine to riboflavin. When we quantified the levels of three flavin end products of the pathway, riboflavin, flavin mononucleotide (FMN) and flavin adenine dinucleotide (FAD) (Fig. 5C, inset), we noted that FMN, the flavin cofactor in FLD1, is the most dramatically increased: over six-fold in the Fe-deficient versus Fe-replete condition. Taken together, these results show a coordinated effort by the cell to replace FDX1 and its 2Fe2S cluster with FLD1 and FMN when the available Fe is low.

### Down regulation of Photosystem I in favor of Photosystem II allows maintenance of photosynthesis with a lower Fe quota

Many of the key components of the photosynthetic apparatus require Fe to function. Since obligately photoautotrophic organisms like *Dunaliella* depend on photosynthesis for life, insufficient Fe can be fatal without countermeasures. About half the Fe in the photosynthetic apparatus is found in photosystem I (PSI). A common acclimation strategy in cyanobacteria is a change in the ratio of PSI to PSII in low Fe to favor PSII (36). We annotated PSI and PSII components in both *D. tertiolecta* and *D. salina* Bardawil and quantified the expression of these genes +/– Fe. All PS genes were very highly expressed, nearly 10^5^ FPKMs for *PSBR1* in *D. tertiolecta*, and all were down regulated in low Fe (SI Appendix, Fig. S8). However, not all PS components were down regulated equally. In *D. tertiolecta*, a PSI component, *PSAE1*, was down regulated 8.7-fold, while a PSII component, *PSBY1*, was down regulated only 1.4-fold. When the down regulation of either PSI or PSII components was averaged, the PSI components were significantly more down regulated (Fig. 6A), consistent with prioritization of PSII over PSI. In *D. salina* Bardawil, the mean log_2_-transformed fold decrease was 1.5 for PSI versus 0.8 for PSII. While the difference was significant in both species, it was more pronounced in *D. tertiolecta*: 2.5 versus 1.0.

**Fig. 6.**
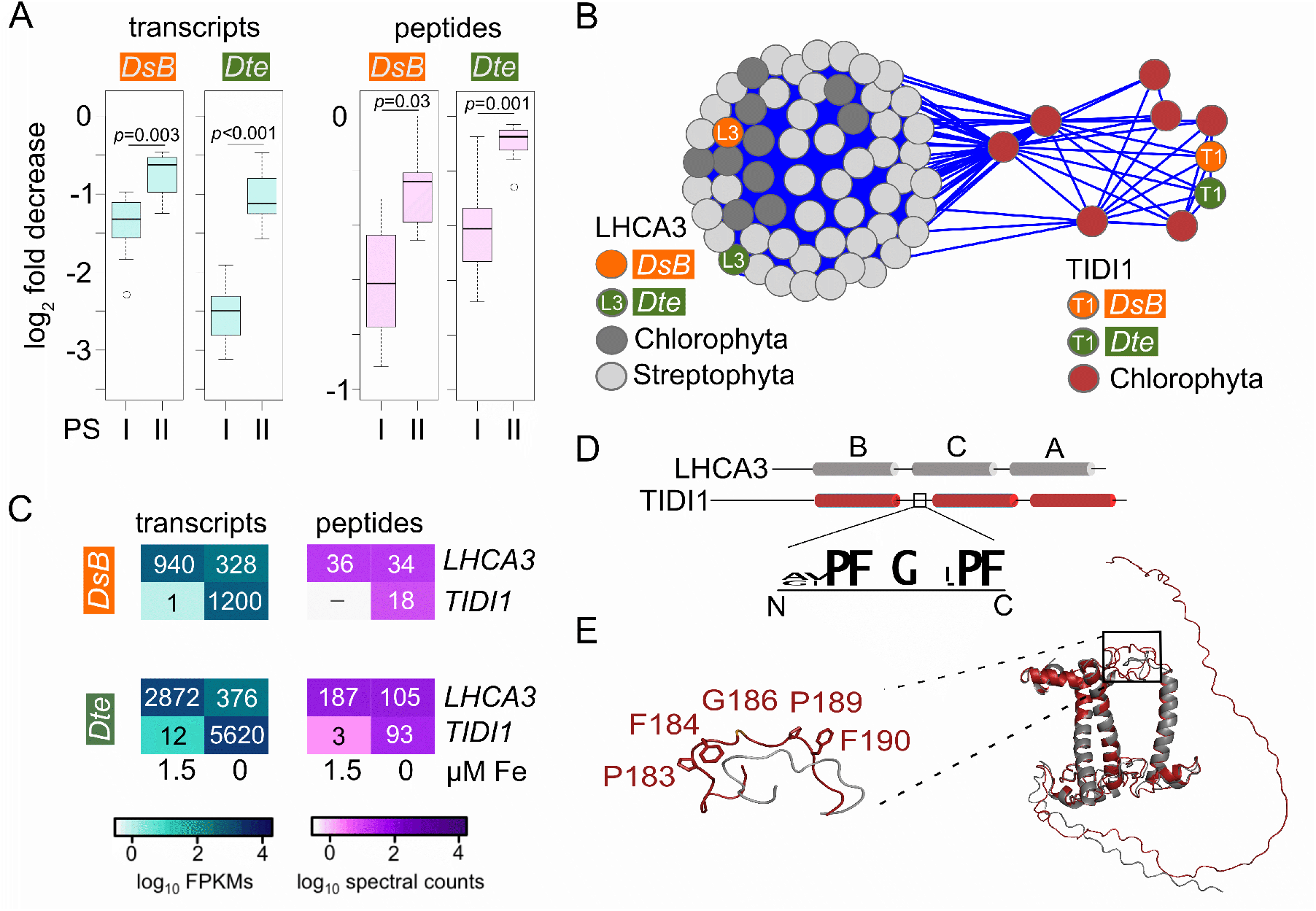
*Dunaliella* remodels its photosystem in response to low Fe, including up regulation of TIDI1. *Dunaliella* species can alter the composition of their photosynthetic machinery to acclimate to low Fe. (**A**) On the left, box plots of the log_2_ fold change decrease in the –Fe versus +Fe condition for transcripts encoding components of PSI (*n* = 9) and PSII (*n* = 7) for *D. salina* Bardawil and *D. tertiolecta*. On the right, the same analysis for PSI (*n* = 7) and PSII (*n* = 7) proteins detected by at least two spectral counts. See also SI Appendix, Fig. S7. (**B**) A protein sequence similarity network of LHCA3 and TIDI1 proteins from Chlorophytes and Streptophytes was generated using an E-value of 1 × 10^−78^. (**C)** Labeled heatmaps of transcript and protein abundance for the products of the *LHCA3* and *TIDI1* genes in *D. salina* Bardawil and *D. tertiolecta* grown in media +/– Fe. (**D**) A cartoon depicts the three alpha helix structure shared by LHCA3 and TIDI1. Below, a logo plot of the conserved amino acids in a loop between alpha helices B and C that is present in TIDI1 but not LHCA3. (**E**) Predicted 3D structures for *D. tertiolecta* LHCA3 (gray) and TIDI1 (red) are overlaid. The black box highlights the location of the 10 aa TIDI1-specific loop and the positions of the most conserved residues.

In previous work, an LHCA3-like protein called TIDI1, expressed exclusively under Fe-starvation conditions, had been discovered in *D. tertiolecta* (reported in that work as *D. salina*) and purported to be an accessory antenna to PSI (14). Indeed, *TIDI1* is highly up-regulated in both *Dunaliella* species, in contrast to canonical *LHCA3* (Fig. 6C). *TIDI1* is not present in *Chl. reinhardtii (*reference alga in the green lineage) or in land plants. However, we discovered TIDI1-encoding genes in the genomes of four other green algae from the order Sphaeropleales: *Chromochloris zofingiensis, Flechtneria rotunda, Tetradesmus deserticola* and *Scenedesmus* sp. NREL_46B-D3. Since *Dunaliella* belongs in order Chlamydomonadales, this suggests that *TIDI1* first arose over 651 mya (Fig. 1B), or else is the result of horizontal gene transfer between species in the Chlamydomonadales and Sphaeropleales lineages. A protein similarity network (Fig. 6B) demonstrates that the Chlorophycean TIDI1 proteins are similar to, but distinct from the LHCA3 proteins of Chlorophytes and Streptophytes.

To explore the relationship between TIDI1 and LHCA3 further, we computationally predicted structures for both proteins and found that while they are nearly identical, two features distinguish them (Fig. 6E). First, the TIDI1 proteins have a longer, proline-rich N-terminal extension relative to LHCA3. Second, TIDI1 proteins have an additional 10-12 amino acids in the loop between alpha helices B and C (Fig. 6D-E). In all the TIDI1 proteins we identified, we found five highly conserved residues in the inter-helix loop arranged as PFxGxxPF. We propose that this pattern could be used as a marker for the identification of TIDI1 as a distinct protein from LHCA3 in other organisms as their genomes are sequenced. In the predicted structure, the orientation of the inter-helix loop places two bulky, hydrophobic Phe residues outward facing (Fig. 6E), which could suggest a TIDI1-specific docking site for binding to the PSI complex.

### *Dunaliella* shares some Fe-related proteins with plants and others with diatoms

There is a rich literature on components of Fe homeostasis in the green lineage and diatoms because of the high abundance of Fe-containing proteins in the photosynthetic apparatus (37–39). For a phylogenomic profile of Fe-homeostasis proteins, we identified orthologs of Fe-related proteins from three Chlorophytes (*D. salina* Bardawil, *D. tertiolecta, Chl. reinhardtii*), two Streptophytes (*Arabidopsis thaliana* and *Oryza sativa*), and three diatoms (*Fragilariopsis cylindrus, P. tricornutum*, and *Thalassiosira oceanica*) (Fig. 7).

**Fig. 7.**
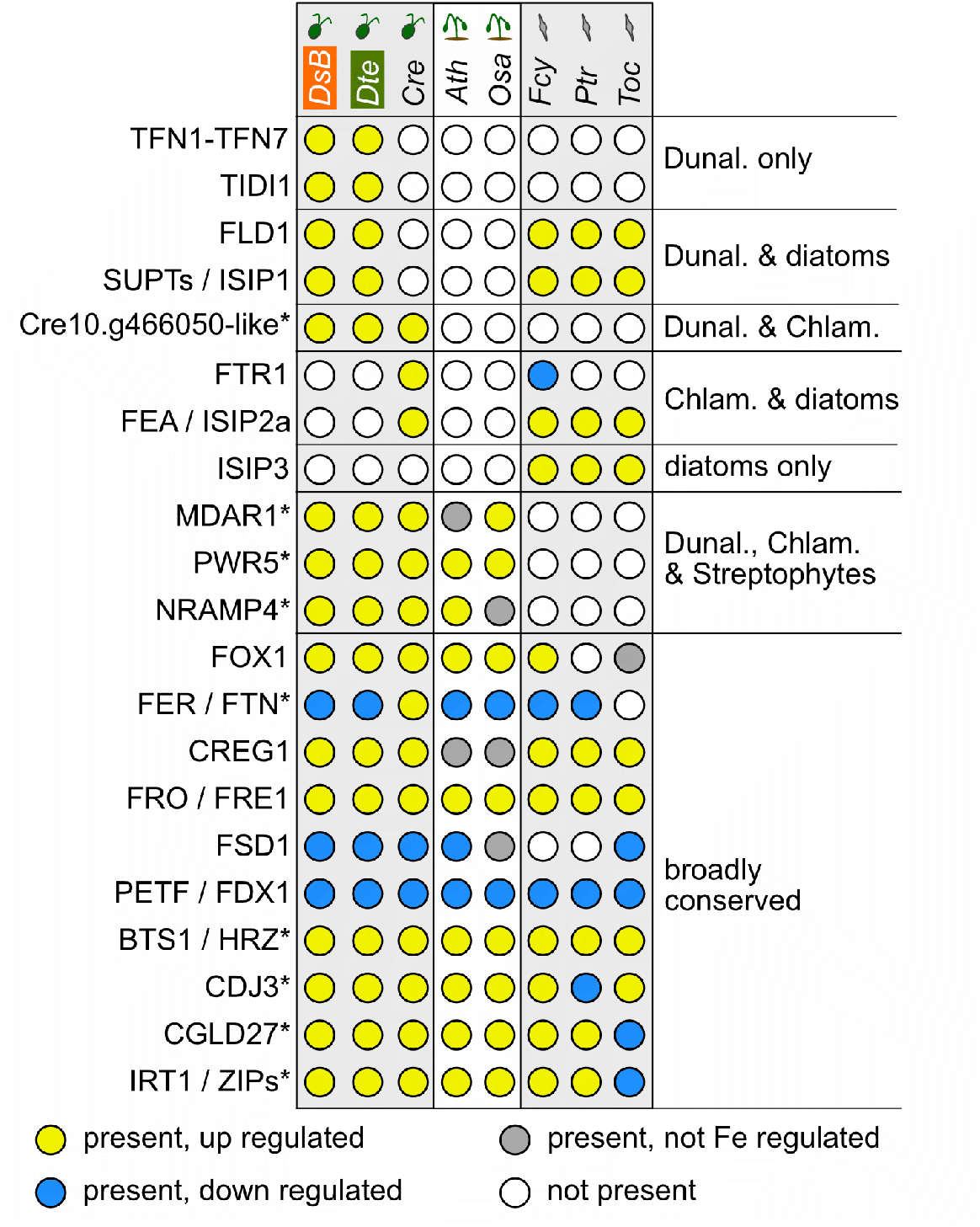
Some Fe-related proteins are conserved among eight photosynthetic species, others are not. Fe-related proteins that were identified in this work and Urzica et al. 2012 (72) were evaluated to identify homologous proteins in eight species of Chlorophytes, Streptophytes and diatoms with RNA-Seq expression data from experiments performed in low and normal Fe. A threshold |log_2_-transformed fold change| > 0.3 was used to define up and down regulation. Species are as follows: Chlorophytes (DsB = *D. salina* Bardawil, Dte = *D. tertiolecta*, Cre = *Chl. reinhardtii)*; Streptophytes (Ath = *Arabidopsis thaliana*, Osa = *Oryza sativa)*; and Diatoms (Fcy = *Fragilariopsis cylindrus*, Ptr = *Phaeodactylum tricornutum*, Toc = *Thalassiosira oceanica)*. Proteins identified as being conserved in the green lineage in (72) are highlighted with an asterisk.

Among these eight species, TIDI1 and the expanded family of Tf proteins are unique to *D. tertiolecta* and *D. salina* Bardawil. Other responses to poor Fe nutrition are conserved between *Dunaliella* and the other members of the green lineage, such as increased expression of monodehydroascorbate reductase 1 (MDAR1) and natural resistance-associated macrophage protein 4 (NRAMP4). In contrast, FLD1 and the SUPT/ISIP1-family proteins are shared only with diatoms, which lie on a different branch of the tree of life, separated from *Dunaliella* by approximately 1.5 billion years of evolution (40).

Recently, the Fe-uptake protein, ISIP2a, was identified in diatoms undergoing Fe insufficiency (41). This protein is sometimes described as phytotransferrin. While this protein may share a common function with the canonical Tf proteins from *Dunaliella*, metazoans and some land plants, it is distinctly different in terms of amino acid similarity and predicted structure (SI Appendix, Fig. S6); likely the result of convergent evolution. Here, we group ISIP2a with the *Chl. reinhardtii* FEA proteins to which they are related (Fig. 7).

Lastly, we note that the two *Dunaliella* species carry orthologs of the *Chl. reinhardtii* gene Cre10.g466050. The protein it encodes has no known function, but its conservation and degree of Fe-regulated expression suggest this is may represent a pioneer protein worthy of further study.

## Discussion

*Dunaliella’s* ability to grow in harsh conditions where the bioavailable Fe is so low as to be growth limiting for neighboring species make it an attractive model for study. When we grew *D. salina* Bardawil cultures in media without added Fe, we found that they were not affected in growth rate or chlorophyll content and were only mildly chlorotic, despite an approximately three-fold decrease in internal Fe content (SI Appendix, Fig. S1). This is in contrast to other algae and land plants that are growth impacted and severely chlorotic under similar conditions. To understand the molecular mechanisms that allow *Dunaliella* to maintain its growth despite Fe scarcity, we produced transcriptomic and proteomic datasets, coupled with biochemical analyses, which facilitated the identification of dozens of genes that contribute to Fe homeostasis in *Dunaliella*.

The range of Fe-related responses that we identified was striking when considering that most other organisms rely on one major strategy for Fe acquisition. Instead *Dunaliella* utilizes multiple, complementary strategies for Fe acquisition (Fig. 8). We describe this as the “mix-and-match” strategy for Fe uptake. First, the reductive Fe-uptake pathway, common in land plants, green algae, and fungi, is characterized by a high uptake rate, but a relatively low affinity for Fe. We observed Fe-dependent expression of the key genes in that pathway, such as *FOX1* and *FRE1A* (Fig. 2). Second, siderophore-mediated Fe uptake, commonly found in bacteria and cyanobacteria, has a high affinity for Fe, but a low rate of uptake. We propose that a family of highly expressed *Dunaliella* proteins of unknown function (previously known as p130B, and here called SUPTs) is used by the cell for siderophore-bound Fe uptake (Fig. 4). And third, Tf-mediated Fe uptake, first identified and commonly found in metazoans, is capable of very high affinity and fast binding of Fe, but requires the production of large quantities of Tf protein, which is energetically costly. We found that both *Dunaliella* species produce several, multi-domain, membrane-bound Tf proteins capable of binding up to 10 atoms of Fe simultaneously (Fig. 3). This major expansion of Tf-encoding genes highlights the importance of this particular strategy for *Dunaliella*. Each of these three different Fe-uptake strategies was first identified in far-flung clades of the tree of life, so it is interesting to consider how the co-occurrence of all three in one species may give *Dunaliella* a fitness advantage.

**Fig. 8.**
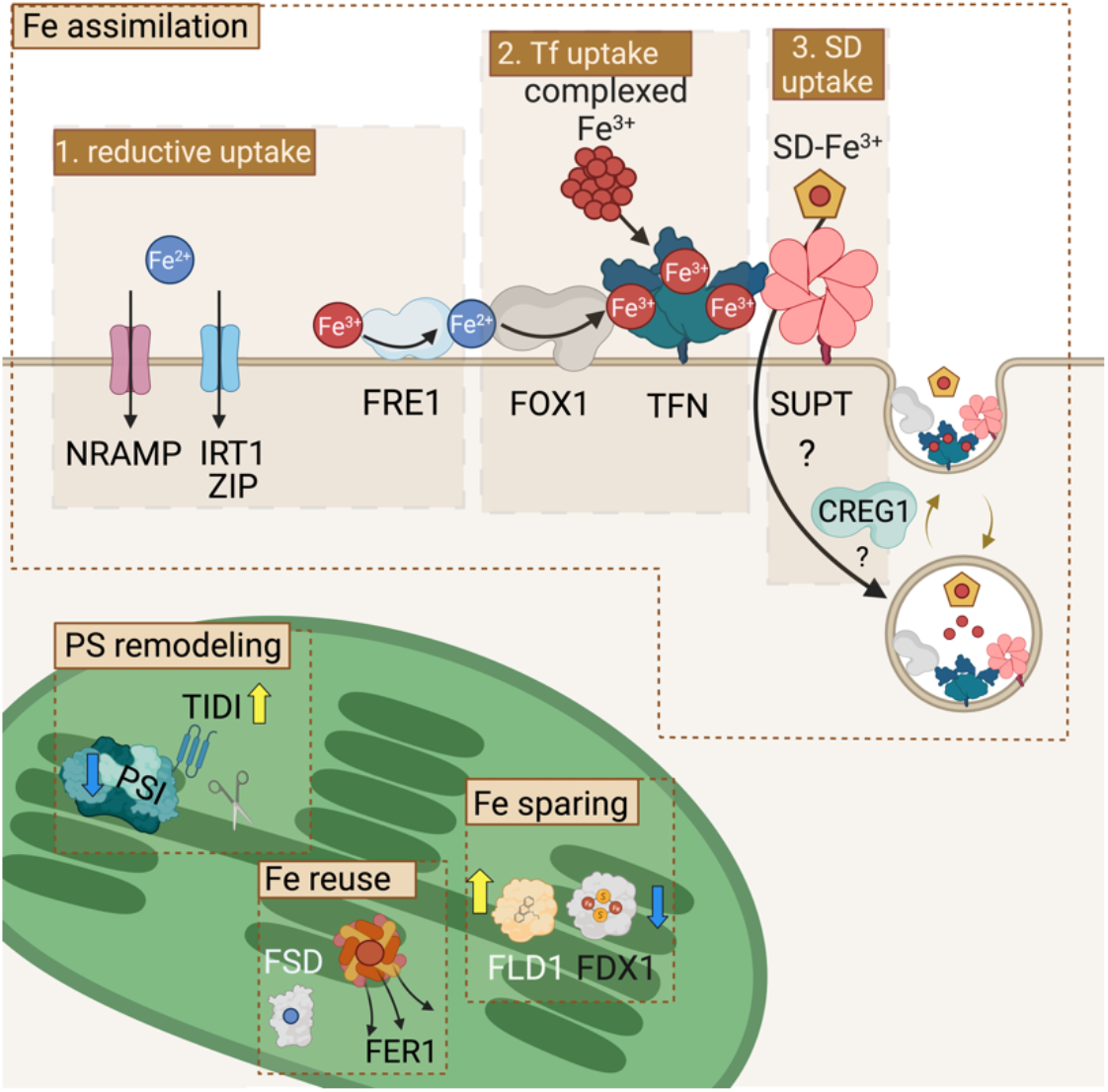
Summary of Fe-responsive mechanisms in *Dunaliella* species. Increased Fe assimilation in *Dunaliella* is mediated by three mechanisms that allow uptake of all forms of Fe. 1. The reductive mechanism, which includes induction of the FRE1A reductase and non-specific divalent metal transporters, like Iron responsive transporter 1 (IRT1), other Zrt/Irt-like protein (ZIP) family transporters, and NRAMP4. 2. Tf-mediated uptake is conferred by a large family of high affinity Fe^+3^-binding proteins. A copper-dependent FOX1, which mediates the uptake of Fe^+2^, may participate in reductive uptake but has also been observed in association with TFN1 (19). 3. Proposed uptake of siderophore-bound Fe via binding to SUPT family proteins. Cellular Repressor of E1A Stimulated Genes 1 (CREG1) is an Fe-regulated protein that may play a regulatory role in Fe uptake. In addition to increased uptake, the Fe quota is maintained by the remodeling of the photosynthetic apparatus, which includes the preferential down regulation of PSI versus PSII components, and the addition of the accessory antenna protein TIDI1. Additional Fe is made available for re-use by degradation of the dispensable Fe superoxide dismutase (FSD) protein and release of Fe stored in FER. Fe sparing occurs by replacing the Fe-rich FDX1A protein with FLD1.

While each of these strategies may function independently, it is interesting to consider the alternative where they work in tandem. A previous study identified components of all three Fe-uptake strategies (TFN1, TFN2, FOX1, and SUPT1) forming a super complex on the plasma membrane of *Dunaliella* (19). Such a super complex would have the ability bind Fe in whichever form it is encountered: Fe^2+^, Fe^3+^ and siderophore-bound. In this model, *Dunaliella* could channel Fe from all three pathways to Tf as follows: from the redox pathway, through FRE1 and FOX1, through the siderophore pathway via the SUPT proteins, and by direct binding to Tf (Fig. 8). In this way, the relative strengths and weaknesses of the three strategies could combine to confer rapid, high-capacity, high affinity, Fe uptake of whichever form of Fe is available.

It is striking how these two *Dunaliella* species have Fe-homeostasis genes in common with Chlorophytes, Streptophytes, diatoms and cyanobacteria. While much of this is certainly due to shared ancestry, we must also consider the possibility that some of these genes are present in *Dunaliella* as the result of horizontal gene transfer. The *SUPT* genes, which we identified in only a few other Chlorophytes, represent an intriguing candidate for further study in this regard.

The high-quality genome assemblies, annotated gene catalogs, and transcriptomics/proteomics datasets that accompany this work should be a vital resource for researchers in a variety of fields. For those engaged in engineering *Dunaliella* species for the production of biofuels and other valuable bioproducts, this work will accelerate those efforts. For those concerned about the role that marine algae can play in climate change mitigation efforts by increasing primary productivity, this work identifies a variety of ways that algae may maintain robust photosynthetic activity despite low available Fe, such as is found in HNLC regions. Finally, the proteins of these *Dunaliella* species are highly evolved to remain correctly folded and functional in up to 3 M NaCl. Just as the thermostable polymerase of *Thermus aquaticus* facilitated the development of PCR, the proteins of *Dunaliella* that we have identified may prove to be an important source of highly-stable proteins for novel applications in synthetic biology and bioengineering.

## Materials and methods

### Strains and culture conditions

*D. salina* Bardawil and *D. tertiolecta* were gifts of Uri Pick. *D. salina* Bardawil is deposited in the University of Texas-Austin Culture Collection of Algae (UTEX) as *Dunaliella bardawil* LB 2538. *D. tertiolecta* is as UTEX LB 999. The medium composition and growth conditions were as described (42). Briefly, all culture glassware were washed in 6 N HCl and thoroughly rinsed with Milli-Q water. Standard growth medium contained 1 M NaCl (*D. salina* Bardawil) or 2 M NaCl (*D. tertiolecta*), with or without the addition of 1.5 µM FeCl_3_, 6 µM EDTA as indicated. Culture flasks were grown on shaking platforms (140 rpm) at 24°C. Constant light at 50 μE m^−2^ s^−1^ (*D. salina* Bardawil) or 100 μE m^−2^ s^−1 (^*D. tertiolecta*) was provided by cool white fluorescent bulbs (4,100 K) and warm white fluorescent bulbs (3,000 K) in a 2:1 ratio. Cells were counted by the Cellometer automated cell counter (Nexcelom).

### DNA preparation and sequencing

Total DNA was isolated from stationary phase cultures of *D. salina* Bardawil and *D. tertiolecta* as described in (43). Short read DNA sequencing for *D. salina* Bardawil and *D. tertiolecta* was performed on the HiSeq 2500 platform (Illumina) with 150+150 paired end libraries following manufacturer’s protocols. Long read sequencing of *D. salina* Bardawil was performed on the Sequel II platform (Pacific Biosciences (PacBio)). Long read sequencing of *D. tertiolecta* was performed on the MinION platform using an LSK-109 library ligation kit and R9 flow cells (Oxford Nanopore Technologies (ONT)). Chromosome Conformation Capture (Hi-C) sequencing was performed on *D. salina* Bardawil by Dovetail Genomics.

### RNA preparation and sequencing

Flasks were inoculated in triplicate at 4 × 10^5^ cells/mL into media with or without 1.5 µM Fe and grown for 2 days (*D. tertiolecta*) or 3 days (*D. salina* Bardawil) under conditions as described above. Cells were collected by centrifugation at 3,500 x *g* for 5 min. Total RNA was purified with TRI Reagent (Molecular Research Center) following manufacture’s protocol. DNA was removed by incubation with TURBO DNase (Invitrogen) following the manufacturer’s protocol. RNA quality was assessed with the RNA Nano kit on a 2100 Bioanalyzer (Agilent).

Short read RNA-Seq for both species was performed on the HiSeq 2500 platform (Illumina) with 50 nt single end libraries following manufacturer’s protocols. A pool of samples of total RNA from *D. salina* Bardawil was used to generate libraries for isomer sequencing (Iso-Seq) on the Sequel II platform (PacBio). Two libraries were prepared and sequenced: one with size-based selection for mRNA >4 kb using AMPure beads (Beckman) and a second library with no size selection.

### Genome assembly

Genome assemblies were generated using a two-step approach: long reads were error corrected with short reads, and the resulting corrected reads were used for genome assembly. In the first step, Illumina short reads were trimmed to remove adaptor sequence with Trimmomatic (v0.39, available at https://github.com/usadellab/Trimmomatic), and coverage-depth normalized to 100x with bbnorm.sh (v38.00, part of the BBMap package available at https://sourceforge.net/projects/bbmap/) using default parameters. Next, the trimmed and down-sampled short reads were used to generate an assembly with MaSuRCa (v3.4.2, available at https://github.com/alekseyzimin/masurca). PacBio long reads from *D. salina* Bardawil and ONT long reads from *D. tertiolecta* were error corrected and trimmed with Canu (v1.8, available at https://github.com/marbl/canu/releases) using the -correct and -trim stages (44). Next, long reads were error corrected with halc (available at https://github.com/lanl001/halc) using three inputs: 1) trimmed and normalized short reads, 2) trimmed long reads, and 3) the short read-only draft assembly from MaSuRCa.

In the second step, the error-corrected long reads were assembled with flye (v2.8.3-b1695, available at https://github.com/fenderglass/Flye) using default parameters (45). Illumina short reads were used for one round of assembly polishing with pilon (v1.22, available at https://github.com/broadinstitute/pilon). Scaffolding of the *D. salina* Bardawil assembly to chromosome scale was performed using Illumina Hi-C reads with Juicer (v1.5.6) and Juicebox (v1.11.08) (both available at https://github.com/aidenlab/Juicebox).

Contigs representing the plastome and mitogenome were identified via sequence similarity to the corresponding organelle genomes from *Chl. reinhardtii* (46) using blastn (47). Annotation of the organelle genomes and generation of maps was performed with OGDraw (https://chlorobox.mpimp-golm.mpg.de/OGDraw.html) (48).

### TEs and repetitive sequence

Preliminary repeat libraries were produced by running RepeatModeler (v2.0.2, available at https://github.com/Dfam-consortium/RepeatModeler) (49) on each genome assembly with the additional LTR structural annotation option (“-LTRStruct”). Consensus repeat models were initially classified by blastx (47) query against a database of TE proteins downloaded from Repbase (50) supplemented with curated proteins extracted from *Chl. reinhardtii* TEs (51). The density of each repeat model was estimated using RepeatMasker (v4.1.2-p1, available at http://www.repeatmasker.org). For repeats contributing >1 Mb of sequence, manually curated consensus sequences were produced following the annotation method of Goubert et al. (52), and seed alignments for upload to the Dfam repository were produced following Storer et al. (53). Additional tandem repeats were annotated using Tandem Repeats Finder (v4.0.9, available at https://tandem.bu.edu/trf/trf.html) (54) with the parameters “2 7 7 80 10 50 2000 -f -d -m -ngs” (i.e. a minimum alignment score of 50 and a maximum monomer length of 2,000 bp). Microsatellites were defined as tandem repeats with monomers <10 bp, with larger monomer repeats defined as satellites. Both classes were supplemented with microsatellites and satellites previously annotated by RepeatMasker. In cases of overlapping microsatellite and satellite annotation, satellites were given precedence since shorter monomers are frequently present within longer repeats. For determining total repeat content, regions annotated as both a TE and tandem repeat were called as TEs. Tandem repeats were given precedence over unknown repeats.

### Gene annotation

Gene annotation was performed using Funannotate (v1.7.2, available at https://github.com/nextgenusfs/funannotate) with default options except --optimize_augustus --species_other --max_intronlen 50000. For *D. salina* Bardawil, the annotation was guided by mapping of 50 nt single-end short RNA-seq reads and long Iso-Seq reads that were generated as part of this study. For *D. tertiolecta*, publicly available 150+150 nt paired-end RNA-Seq reads (55) and 150 nt single-end reads (56) were used as inputs, along with the Funannotate-predicted proteome of *D. salina* Bardawil. Gene loci were assigned IDs of the form “DsB_gXXXXX” for *D. salina* Bardawil and “Dte_gXXXXX” for *D. tertiolecta*, where “XXXXX” indicates a five-digit serial number.

RNA-Seq reads from *D. salina* Bardawil and *D. tertiolecta* were mapped to their respective genome assemblies with STAR (v2.4.0j, available at https://github.com/alexdobin/STAR) with the --alignIntronMax 5000 flag. Iso-Seq reads from *D. salina* Bardawil were mapped with minimap2 (v2.17-r941, available at https://github.com/lh3/minimap2) with -a -x splice. Gene models were extensively reviewed and manually edited as needed base on the alignment of RNA-Seq and Iso-Seq reads as viewed using the IGV browser (v2.9.4, available at https://software.broadinstitute.org/software/igv/) (57).

Gene symbols were assigned based on extensive manual curation. Additional gene symbols were assigned based on orthology relationships with *Chl. reinhardtii* v6.1 (58) as identified with orthofinder (v2.5.2, available at https://github.com/davidemms/OrthoFinder).

Exon and intron sizes and their distributions were calculated with calculate_exon_intron_sizes.pl (v1.0, available at https://github.com/seangallaher/calculate_exon_intron_sizes).

### RNA-Seq analysis

RNA-Seq reads were aligned with STAR as described above. Normalized expression estimates in terms of fragments per kb of sequence per million mapped reads (FPKMs) were calculated using cuffdiff (v2.0.2, available at http://cole-trapnell-lab.github.io/cufflinks/) with the following flags: --multi-read-correct --max-bundle-frags 1000000000 --library-type fr-firststrand. DEGs were determined with DESeq2 (59). DEGs were defined as having a |log_2_ fold-change| >1 and the Benjamini-Hochberg adjusted *p*-value <0.01.

### BUSCO analysis

Genome and transcriptome quality were assessed using BUSCO (v4.0.5, available at https://busco.ezlab.org/) using the chlorophyta odb10 dataset (60).

### Phylogenies and Chronograms

For the Chlorophyte species chronogram, the proteomes of 12 Chlorophycean algae were subjected to BUSCO analysis as described above. Those proteins that were present as single copy orthologs in all 12 species were isolated (104 total) and subjected to multiple sequence alignment with Muscle (v3.8.31, available at http://www.drive5.com/muscle/) using default parameters (61), followed by end trimming with trimAl (v1.2rev59, available at http://trimal.cgenomics.org) (62), and concatenation of the trimmed sequences. For the transferrin chronogram, candidate proteins were collected from public databases of proteins by sequence similarity and/or presence of the transferrin PFAM domain (PF00405). Polypeptide sequences were subjected to multiple sequence alignment with Muscle, and retained or rejected based on the presence of conserved Fe and anion binding residues (18).

Phylogenetic trees were generated by Bayesian Monte Carlo Markov Chain sampling with phylobayes (v4.1c, available at http://www.atgc-montpellier.fr/phylobayes/) (63). Specifically, the pb algorithm was run with two parallel chains until the maxdiff < 0.3 as determined by bpcomp -x 100 2. The majority-rule posterior consensus tree was generated with readpb -x 100 10. Molecular dating was estimated with the phylobayes pb algorithm using an uncorrelated gamma multiplier relaxed clock model (-ugam). For the species chronogram, calibrations were from Del Cortona et al. (6), and Trebouxiophyceaen sequences were used as an outgroup. For the transferrin chronogram, calibrations were from TimeTree (http://timetree.org/) (40), and Metazoan transferrin sequences were used as an outgroup. Chronogram figures were constructed using the Interactive Tree of Life webapp (https://itol.embl.de/) (64).

### Protein structure predictions

Protein structure predictions were conducted using ColabFold (65) (https://github.com/sokrypton/ColabFold). Structures were visualized with PyMOL (v1.7.4, available at https://pymol.org/). Logo plots were generated with WebLogo (available at https://weblogo.berkeley.edu/) (66).

### Orthology analysis and estimation of gene duplication events

Orthological and paralogical relationships between proteins, and estimation of gene duplication events were determined by OrthoFinder (v2.5.2, available at https://github.com/davidemms/OrthoFinder) using default parameters (67). A full list of the species used, their version numbers, and their sources is provided in SI Appendix, Table S3.

### Protein similarity networks

Protein similarity networks were generated from an all-versus-all blastp (47) analysis (pairwise alignment between all pairs of proteins) of sequences in a local sequence database. The networks were created in Cytoscape (v3.4) with the BLAST2SimilarityGraph plug-in and the yFiles Organic layout engine provided with Cytoscape. Protein nodes were connected by edges if the E-value between the two sequences was at least as good as the value indicated the the corresponding figure legend.

### Synteny analysis

Synteny between the *D. salina* Bardawil and *D. tertiolecta* assemblies was determined with SynChro (vJanuary 2015, available at http://www.lcqb.upmc.fr/CHROnicle/SynChro.html) (68), and visualized with Circos (v0.69-9, available at http://circos.ca/) (69).

### Proteomics and SRM quantification

Total soluble protein was collected in parallel with the RNA-Seq samples described above, tryptic digested, and quantified by mass spectrometry as described previously ((46). For SRM quantification of FLD1 and FDX1, peptides were selected that were readily detectable in other samples and that could be uniquely assigned to their respective protein. Heavy versions of these peptides were synthesized with ^13^C- and ^15^N-enriched amino acids to label the terminal K or R. Two-fold dilution series of each heavy peptide were added to cellular protein samples and assayed to validate the linear range of detection. The optimal concentration of each heavy peptide was added to biological samples in triplicate and assayed. The ratio of light peptide to the spiked-in quantity of the heavy peptide was then used to calculate the quantity of the light peptide on a per cell basis. Three peptide sequences were used for FLD1: QYDGLIVGSPTWNTGADEER, WAYSEGEYEHTYSK and VDKWVAQIR. One peptide sequence was used for FDX1A: VESGTVDQSDQSFLDDDQQGR.

### Elemental composition

Cells were cultured as indicated above. At indicated times, 5 × 10^7^ cells from each flask were collected by centrifugation at 3,500 ×*g* for 5 min and washed three times with 1 M NaCl and once with Milli-Q water. After removing the remaining water by a brief centrifugation, cell pellets were digested with 70% nitric acid at 65°C for 2 h. Digested samples were diluted with Milli-Q water to a final nitric acid concentration of 2% (v/v). The elemental composition was measured by ICP-MS as described in (70).

### Metabolite analysis

*D. salina* Bardawil cultures were grown in media +/– Fe as described above. On day 3, cells were counted and 50 mL from each flask were collected by centrifugation at 3,500 x *g* for 5 min and washed three times with 1 M NaCl. Cell pellets were weighed and flash frozen in liquid nitrogen. Metabolites were analyzed as described previously (71).

### Chlorophyll content

Chlorophyll content was assayed as described previously (42).

### Statistical analyses

All *n* numbers represent biological replicates (i.e. samples taken from independent culture flasks). Data in bar graphs are expressed as the mean ± standard deviation. Significance was determined by two-sided Student’s t-test. For boxplots, the center line indicates the median, box limits indicate the upper and lower quartiles, whiskers indicate 1.5x the interquartile range, and points indicate outliers.

## Code availability

Custom code for calculating exon and introns sizes from a transcriptome is freely available from Github under a GNU General Public License at https://doi.org/10.5281/zenodo.7719065.

## Data availability

Genomic sequencing data were submitted to the NCBI Sequence Read Archive (SRA) under BioProject PRJNA914849 (reviewer link: https://dataview.ncbi.nlm.nih.gov/object/PRJNA914849?reviewer=c307nedjc29p85kqbmjivf7iq4) for *D. salina* Bardawil and BioProject PRJNA914763 (reviewer link: https://dataview.ncbi.nlm.nih.gov/object/PRJNA914763?reviewer=vdd273tjrps2heh48ht14vvcbb) for *D. tertiolecta*. Transcriptomic data, assembled genomes, and annotation files were submitted to NCBI Gene Expression Omnibus (GEO) archive under accession GSE222140 (token for reviewer access = wjqzoyckftqvvqr) for *D. salina* Bardawil and GSE222141 (token for reviewer access = kzabawqsjrcrlkd) for *D. tertiolecta*. Accession codes for all other data not generated as part of this work are detailed in SI Appendix, Table S3. Expression data generated in this work are provided in SI Dataset S1.

## Acknowledgments

We want to thank Uri Pick for his helpful comments. LD was supported by the European Molecular Biology Organization (ALTF 166-2016) and by the United States–Israel Binational Agricultural Research and Development (BARD) Fund, Vaadia-BARD Postdoctoral Fellowship Award FI-531-2015. This work was supported by US Department of Energy (DOE), Office of Science, Office of Basic Energy Sciences Award (DE-FG02-04ER15529). Proteomics analyses were supported by an award (49840) from the Environmental Molecular Sciences Laboratory (EMSL; grid.436923.9), which is a DOE Office of Science User Facility under contract DE-AC05-76RL01830. The research conducted by CEBH at the U.S. Department of Energy Joint Genome Institute (https://ror.org/04xm1d337), a DOE Office of Science User Facility, is supported by the Office of Science of the U.S. Department of Energy operated under Contract No. DE-AC02-05CH11231.

## Notes

### Competing Interest Statement

The authors have declared no competing interest.

### Summary of Updates

Minor update.

