## Supplemental Information for "Pumping Iron: A Multi-omics Analysis of Two Extremophilic Algae Reveals Iron Economy Management"

[1] Department of Chemistry & Biochemistry, University of California, Los Angeles, Los Angeles, CA 90095, USA
[2] California Institute for Quantitative Biosciences (QB3), University of California, Berkeley, Berkeley, CA 94720, USA
[3] Department of Human Genetics, University of California, Los Angeles, Los Angeles, CA 90095, USA
[4] Environmental Molecular Sciences Laboratory, Pacific Northwest National Laboratory, Richland, WA 99354, USA
[5] Biological Sciences Division, Pacific Northwest National Laboratory, Richland, WA 99354, USA
[6] Institute of Biological Chemistry, Washington State University, Pullman, WA 99163, USA
[7] DOE Joint Genome Institute, Lawrence Berkeley National Laboratory, Berkeley, CA 94720, USA
[8] Molecular Foundry, Lawrence Berkeley National Laboratory, Berkeley, CA 94720, USA
[9] Purdue Center for Plant Biology, Department of Biochemistry, Purdue University, West Lafayette, IN 47907, USA
[10] Department of Plant and Microbial Biology, Department of Molecular and Cell Biology, University of California, Berkeley, Berkeley, CA 99354, USA
[11] These authors contributed equally.
[12] Present address: Illumina Artificial Intelligence Laboratory, Illumina, Inc., Foster City, CA 94404, USA
[13] Present address: Plant Research Laboratory, Department of Biochemistry and Molecular Biology, Michigan State University, East Lansing, MI 48824, USA

**This PDF file includes:**

Supporting text
Figures S1 to S13
Tables S1 to S6
Legend for Dataset S1
SI References

**Other supporting materials for this manuscript include the following:**

Dataset S1

#### Supporting text

***D. tertiolecta*** **and** ***D. salina*** **Bardawil have exceptionally large genomes.** In this work, we employed a systems biology strategy to elucidate how two *Dunaliella* species are able to thrive in low-Fe environments. This approach requires high-quality genome assemblies to provide a solid foundation for the analysis. Prior to this work, there was no publicly-available genome assembly for *D. tertiolecta*. Two independently produced *de novo* transcriptome assemblies have been published for the species (1, 2), but both have a high degree of gene fragmentation and duplication (Table S4), and were found to be of limited utility. For *D. salina*, a draft genome assembly from strain CCAP 19/18 was published in 2017 (3), but this assembly is also significantly fragmented, limiting its utility (Table S4). For example, the multicopper ferroxidase 1 gene (*FOX1,* previously identified as D-Fox), which was of interest for this study, was found to be in fragments scattered over nine separate ~1-3 kb scaffolds in the *D. salina* CCAP 19/18 assembly. The *FOX1* ortholog in *D. tertiolecta* was completely missing from that species' *de novo* transcriptome (2).

Given the limitations of the previously available assemblies, we opted to generate high-quality, *de novo* assemblies for both species. For *D. salina* Bardawil, this effort produced a 363 Mb nuclear assembly (Table S5). 99.1% of which was scaffolded into 16 chromosomes (Fig. S9), with the remaining 0.9% in 193 unplaced scaffolds. Gaps, in the form of Ns, make up 0.11% (384,490 bp) of the assembly. This assembly is slightly bigger than the 344 Mb assembly previously reported for *D. salina* CCAP 18/1973, and substantially less fragmented (Table S4). The N50/L50 of contigs in our *D. salina* Bardawil assembly was 1016 kb/106 contigs, as compared to 7 kb/11,486 contigs for the *D. salina* CCAP 18/19 draft assembly (Table S4).

For *D. tertiolecta*, we produced a 190 Mb nuclear genome assembly. While not fully at chromosome scale, the *D. tertiolecta* nuclear genome assembly was ordered into 18 scaffolds, and is slightly less fragmented than the *D. salina* Bardawil assembly. The *D. tertiolecta* assembly has a contig N50/L50 of 4,336 kb/14 contigs (Table S4). The percentage of gaps in the assembly was similar to the *D. salina* Bardawil assembly: 181,322 Ns representing 0.096% of the assembly. Finally, we observed a large degree of synteny between the two assemblies (Fig. 1C). In our evaluation of the new genome assemblies for *D. tertiolecta* and *D. salina* Bardawil, we noted that both were remarkable for their size compared to other green microalgae, including the reference species *Chl. reinhardtii* (Fig. 1D). At nearly 200 Mb, *D. tertiolecta*'s genome is larger than most sequenced chlorophytes, but the more extremophilic species, *D. salina* Bardawil, is 91% larger still. These observations prompted the question: what does this "extra" genome sequence in the *Dunaliella* genomes consist of? The following sections consider this question.

**The *D. tertiolecta* and *D. salina* Bardawil genome assemblies are rich in TEs and repetitive elements.** One possible explanation for the large genome sizes of *D. salina* Bardawil and *D. tertiolecta* could be an expansion of TEs and other repetitive sequences. To address this, we annotated and quantified the transposable elements (TEs) and tandem repeats (satellites and microsatellites) in both assemblies (Table S1).

The two *Dunaliella* genomes generally contained a common repertoire of TE orders and superfamilies, most of which are also found in *Chl. reinhardtii*. These include three LINE clades, *Dualen*, *RTE-X* and *L1*, and both *Gypsy* and *Copia* LTR retrotransposons (Fig. S2). While *Dualen* LINEs were abundant in all three genomes, *RTE-X* elements contributed far more, and *L1* elements far less, to the *Dunaliella* genomes, relative to *Chl. reinhardtii*. This was partly due to the absence of *Zepp*-like *L1* elements in *Dunaliella*, which likely constitute the centromeric repeats and are the most abundant TE in *Chl. reinhardtii* (4). The *Dunaliella* genomes contained *hAT* and *EnSpm* DNA transposons, as found in *Chl. reinhardtii*, and *Harbinger* transposons, which have not been reported from green algae but are widespread in plant genomes and present in red algae (5). The *Dunaliella* genomes lacked the *Penelope*-like elements and Helitrons characteristic of *Chl. reinhardtii* (6, 7), although DIRS retrotransposons were present at very low copy numbers.

The overall repeat content of *D. tertiolecta* was 45.5% (86.4 Mb), relative to 60.3% (218.6 Mb) in *D. salina* Bardawil (Table S1). Unlike the more compact *Chl. reinhardtii* genome (25.0% repeats, 27.7 Mb), degraded and likely inactive TEs comprised the majority of repeats in both *Dunaliella* genomes (Fig. S2). The greater repeat content of *D. salina* Bardawil could generally be attributed to an increase of both older and younger TEs proportional to its genome size, rather than a recent burst of any particular TE clade. *Dualens*, giant LINEs generally exceeding 10 kb in length (8), were the most abundant TEs in both genomes, spanning 14.6% and 12.8% of the *D. tertiolecta* and *D. salina* Bardawil genomes, respectively. These were followed by *RTE-X* LINEs, which appeared to be undergoing a recent expansion in *D. tertiolecta* and were proportionally more abundant (8.7% of the genome) than in *D. salina* Bardawil (4.4%; Fig. S2). *Gypsy* elements were the most abundant LTRs and were the only group to show a strong signature of recent expansion in *D. salina* Bardawil, although they span only 4.0% of the genome (0.74% in *D. tertiolecta*). DNA transposons, most of which were degraded nonautonomous families, were substantially more abundant in *D. salina* Bardawil (9.9% of the genome) than *D. tertiolecta* (3.6%). Finally, the *D. salina* Bardawil genome contained a larger proportion of tandem repeats (16.3%, relative to 9.2% in *D. tertiolecta*), which was largely driven by a ~3.5x greater abundance of microsatellites in *D. salina* Bardawil relative to genome size.

Overall, repeats contribute substantially, but cannot entirely explain, the differences in genome size between these species: the *D. salina* Bardawil genome features 144.2 Mb of non-repetitive sequence relative to 103.4 Mb in *D. tertiolecta* and 83.3 Mb in *Chl. reinhardtii*. This prompted us to look at the gene annotations to explain the remaining difference in their respective genome sizes.

**Gene annotation of the *D. tertiolecta* and *D. salina* Bardawil assemblies reveals large introns and significant gene expansion.** We annotated the genomes of *D. salina* Bardawil and *D. tertiolecta* with three goals in mind: 1) to identify the source of the remaining "extra" DNA, 2) to form the foundation for a systems biology analysis of Fe homeostasis, and 3) for gene discovery. This process identified 14,351 protein-coding genes in *D. tertiolecta* (Table S6). This number of genes is approximately one-fourth of the 56,926 transcripts in the *de novo* transcriptome published by the Lee group (1). When analyzed with the chlorophyte dataset, the *D. tertiolecta* transcriptome presented in this work represented a major improvement in terms of missing, fragmented, and duplicated BUSCO scores relative to both previously published transcriptomes (Table S4). Additionally, we estimate that 36.7% of the *D. tertiolecta de novo* transcriptome produced by the Cho group (2) is bacterial in origin, likely due to contamination in their cultures.

With approximately 14,000 nuclear protein-coding genes, *D. tertiolecta* is very near the median for transcriptome size in annotated chlorophytes (Fig. 1F). In contrast, we identified 25,074 protein-coding genes in *D. salina* Bardawil, which is well above the number of genes for the other microalgal species examined. This high number of genes is due in part to the proliferation of TEs described above. Analysis of chlorophyte BUSCO scores indicated slightly fewer missing, and slightly more duplicated genes in the transcriptome of *D. salina* Bardawil relative to *D. tertiolecta* (Table S4).

To further evaluate their quality, the gene models in both species were subjected to manual review relative to the RNA-Seq and Iso-Seq data. During this process, we observed that many of the genes in both *Dunaliella* species had extremely long introns relative to other chlorophyte species, such as *Chl. reinhardtii;* an observation that could help explain the large genomes. To quantify this systematically, we computed the size distribution of the coding sequence (CDS) and intronic portions of all genes for both *Dunaliella* species, and two other chlorophyte algae: *Chl. reinhardtii* and *Chr. zofingiensis* (Fig. S3). In all four species, the overall distribution of protein-coding nucleotides per gene was fairly similar, with the CDS of most genes falling well under 5,000 bp. *Chl. reinhardtii* had the largest CDSs on average, with a mean of 2,206 bp, compared to 1,197 bp in *D. salina* Bardawil, 1,553 bp in *D. tertiolecta*, and 1,444 bp in *Chr. zofingiensis* (Fig. S3A). In contrast, the mean of intronic nucleotides per gene was dramatically higher for the two *Dunaliella* species: 6,902 nt and 6,719 nt for *D. salina* Bardawil and *D. tertiolecta*, respectively, compared to 2,006 nt for *Chl. reinhardtii* (Fig. S3B). The fact that *Dunaliella* genes have significantly more intronic nucleotides than those of the other algae examined here is not due to there being more introns per gene, but rather to the individual introns being larger. Both *Chl. reinhardtii* and *D. salina* Bardawil have similar numbers of introns per gene (7.5 and 5.3, respectively) (Fig. S3E). However, the mean *D. salina* Bardawil intron is over 6x larger: 1,213 nt versus 266 nt in *Chl. reinhardtii* (Fig. S3D).
Given that *Dunaliella* species have such large introns, we wished to determine if that feature could make a significant contribution to their exceptionally large genomes. To that end, all nucleotides in the genomes of the two *Dunaliella* species, as well as the assembled genomes of many other species, were categorized as being inter-CDS, CDS, or intronic (Fig. 1E). Both *Dunaliella* species had the largest contribution of intronic nucleotides of any of the species examined, making up approximately half of their respective genome assemblies: 51% for *D. tertiolecta* and 48% for *D. salina* Bardawil.

In the course of functionally annotating the genes of *D. salina* Bardawil and *D. tertiolecta*, we observed a number of large gene families, especially in *D. salina* Bardawil, which suggested there may have been widespread gene expansion. Recently, a study of another extremophilic alga, *Chl. priscuii*, identified gene expansion as a feature of that species (9). To examine this in *Dunaliella*, we computationally predicted gene duplications. In agreement with our anecdotal observations of gene expansion, *D. salina* Bardawil was predicted to have significantly more gene duplications than any other chlorophyte (Fig. 1G).

Thus, multiple factors contribute to the large genome sizes of these two *Dunaliella* species. Namely, an abundance of repetitive sequence that includes many inactive/degraded TEs, significantly larger introns, and an expansion of gene families all contribute to make the *D. salina* Bardawil and *D. tertiolecta* genomes larger than those of mesophilic green algae like *Chl. reinhardtii*.

**The organellar genomes and transcriptomes of *D. salina* Bardawil and *D. tertiolecta***
The organelles make an often overlooked, but significant contribution to the genetic diversity and transcriptomic capacity of eukaryotic organisms like *Dunaliella*. We assembled the genomes of the chloroplast (plastome) and mitochondrion (mitogenome), and annotated their protein-coding genes. Both *Dunaliella* species have a circular plastome, 265 kb of *D. salina* Bardawil (Fig. S10) and 243 kb for *D. tertiolecta* (Fig. S11), which are slightly larger than, but generally similar in structure and number of protein-coding genes, to the plastomes of other chlorophytes like the 206 kb plastome of *Chl. reinhardtii* (10). Structurally, both plastomes have two inverted repeats (IRA and IRB) which are separated by a long single-copy region (LSC) and a short single-copy region. Owing to their origins as bacterial endosymbionts, most genes in plastomes do not have introns, but there are exceptions. In *Chl. reinhardtii*, only the *psbA* gene, which encodes the photosystem II (PSII) D1 protein, has four type-I introns. In contrast, multiple plastid genes in *Dunaliella* have introns, up to eight genes in *D. tertiolecta*: *psbA, petL, psbD, atpA, psbC, atpH, psaB*, and *psaA*. The gene that encodes PSI chlorophyll *a* binding apoprotein, *psaA*, is striking for the complexity of its transcript. Like its ortholog in *Chl. reinhardtii*, the mature *psaA* transcript is formed by trans-splicing three independently transcribed segments. However, unlike *Chl. reinhardtii*, the *psaA* genes of *Dunaliella* also have introns (one in *D. salina* Bardawil and two in *D. tertiolecta*) that must be excised to form the mature transcript. It is remarkable that this vitally important protein has such a complicated structure.

While each cell of *Dunaliella* has a single plastid, each plastid contains many copies of the plastome. As was done for *Chl. reinhardtii* (10), we calculated the relative mean DNA-Seq coverage of the nuclear genome and the plastome to estimate that each cell of *D. salina* Bardawil carries 135 copies (±51) of the plastome. For *D. tertiolecta*, that number is 52 copies (±11) of the plastome per cell.

The mitogenomes of *Dunaliella* are 29 kb for *D. salina* Bardawil (Fig. S12) and 42 kb for *D. tertiolecta* (Fig. S13). While, *Chl. reinhardtii* has a linear mitogenome (10), the mitogenomes of *Dunaliella* are in the more common circular configuration. Like *Chl. reinhardtii*, *Dunaliella* mitogenomes are highly reduced. They carry only seven protein-coding genes, which encode components of complex I, III, and IV. Introns are common in the mitochondrial genes, with five out of seven genes in *D. tertiolecta* bearing introns. Based on DNA-Seq coverage, we estimate that there are 158 copies (±50) of the mitogenome per cell in *D. salina* Bardawil and 65 copies (±17) in *D. tertiolecta*.

**A stop codon in *D. salina* Bardawil *SUPT1* is likely the result of a recent nonsense mutation.** In this work, we identified a family of putative siderophore-Fe uptake proteins (SUPT) in both species of *Dunaliella* (Fig. 4). All five *SUPT* genes were highly up regulated in the Fe-deficient condition at the transcript level, and four of five were also up regulated at the protein level (Fig. 4A). However, the fifth, *D. salina* Bardawil *SUPT1*, was undetected at the protein level. Upon closer examination of that gene, we identified a stop codon in the middle of an otherwise intact 3216 nt ORF (Fig. 4A). If that stop codon were skipped, the resulting gene would encode a 1071 amino acid protein that is highly conserved with the orthologous SUPT1 proteins from *D. tertiolecta* and from *D. salina* CCAP 19/18 (Fig. S7). Given that the rest of this ORF is intact, with comparable amino acid conservation both upstream and downstream of the stop codon, we hypothesize that a single nucleotide variant has introduced a Lys431* nonsense mutation in that gene. The presence of a premature stop codon would then explain the absence of detected peptides for this gene due to nonsense-mediated decay. We speculate based on the low number of variants in this gene relative to its *D. tertiolecta* ortholog, and the absence of any other stop codons, that this mutation may have occurred fairly recently, perhaps in the laboratory isolate. This might be expected for a gene that is not required for growth under laboratory conditions (i.e. in Fe-replete medium). There is precedent for this phenomenon from *Chl. reinhardtii,* where most laboratory strains have accumulated multiple mutations in two nitrate reductase genes that are superfluous for growth in commonly used, ammonium-containing culture media (11).

### Supporting figures

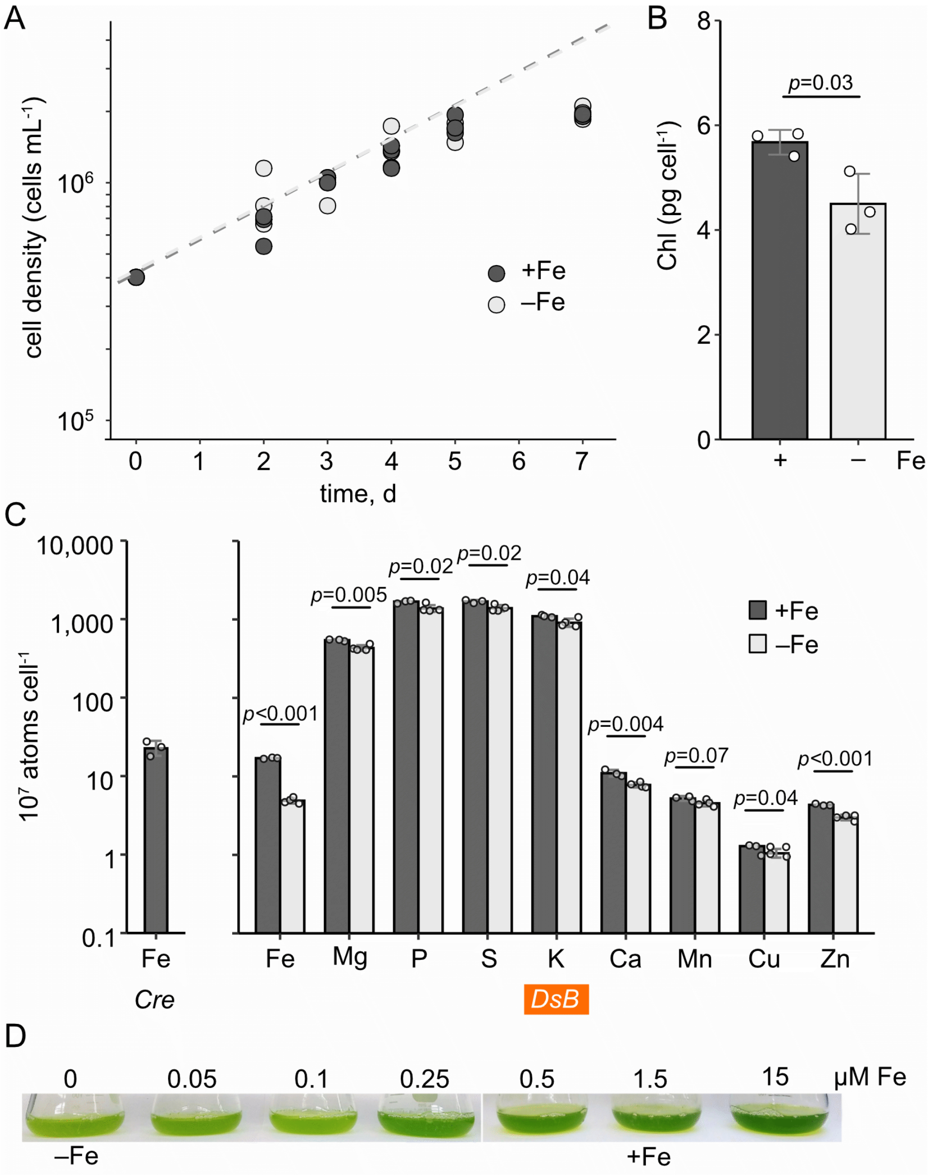


**Fig. S1. Effects of Fe on cell growth, Chlorophyll content and intracellular elemental content.** Cultures of *D. salina* Bardawil were inoculated at 5 x $10^5$ cells/mL into media with (dark gray) and without (light gray) 1.5 µM of added Fe. (**A**) Cell counts were performed from cultures grown ±Fe over seven days of growth and are plotted on a semi-log scale ($n$ = 3). Dashed lines depict linear regressions of the log growth phase timepoints (d0 – d5) for + and –Fe cultures. (**B**) The Chlorophyll (Chl) content was calculated on a per cell basis for samples collected on day 7 ($n$ = 3). (**C**) The intracellular elemental content was determined by ICP-MS and plotted on a per-cell basis for cultures of *D. salina* Bardawil (*DsB*) grown in media +Fe ($n$ = 3) and –Fe ($n$ = 4). The intracellular content of Fe for *Chl. reinhardtii* cultures grown in 20 µM Fe (replete for that species) is shown on the left ($n$ = 3). (**D**) Photographs of culture flasks grown with increasing concentrations of added Fe. Throughout this work, -Fe corresponds to 0 µM and +Fe corresponds to 1.5 µM added Fe as indicated below this figure.

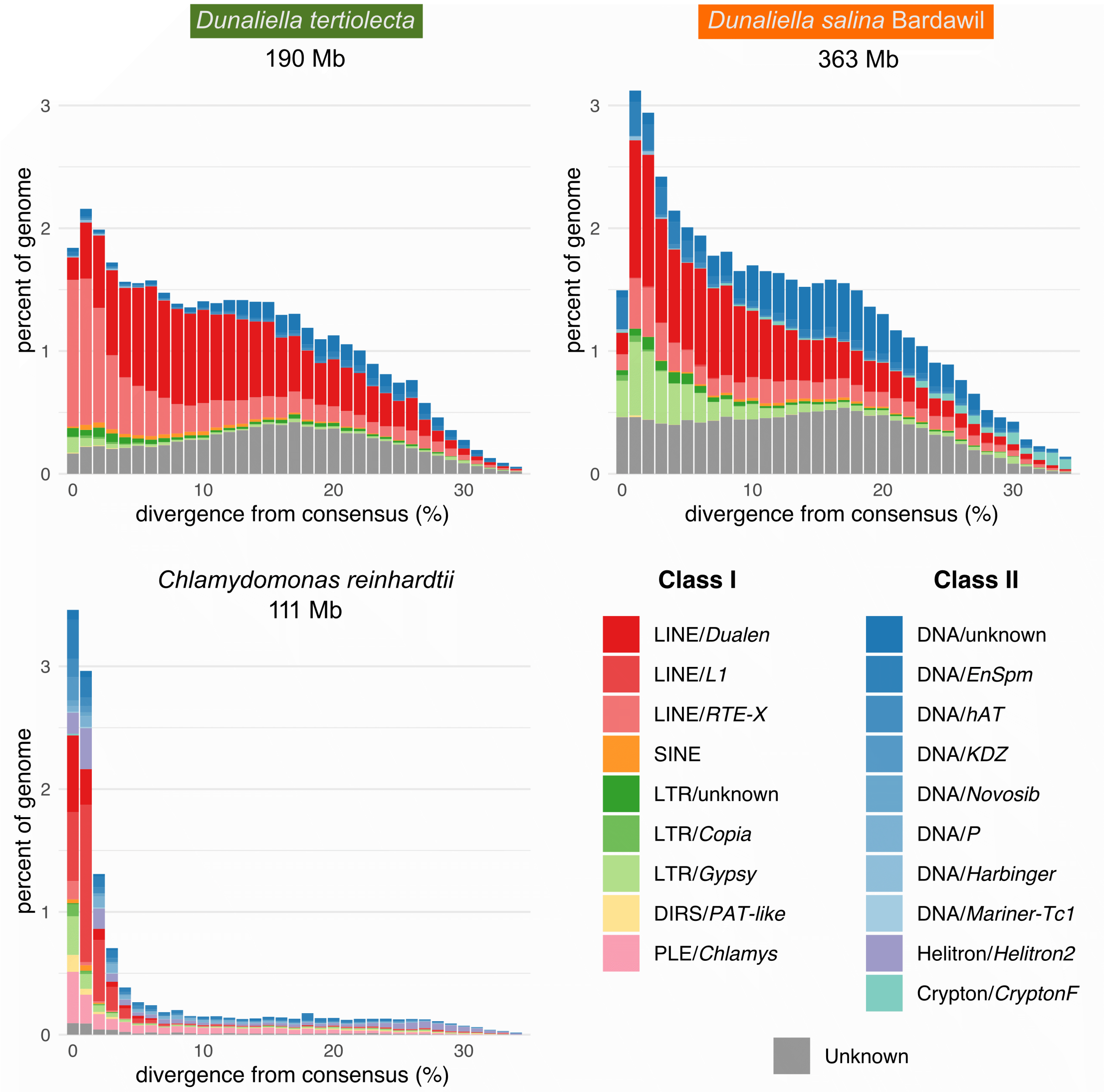


**Fig. S2. Transposable elements and repeat content in the *Dunaliella* and *Chlamydomonas* genome assemblies.** Transposable element landscapes for *D. tertiolecta, D. salina* Bardawil and *Chl. reinhardtii* CC-1690. The nuclear genome size of each species is provided. The divergence of individual TE copies from their consensus repeat model was calculated by RepeatMasker using the Kimura two-parameter model. *Dunaliella* genomes were annotated using their respective hybrid curated/automated repeat libraries (see Methods) and *Chl. reinhardtii* was annotated using a curated repeat library (4).

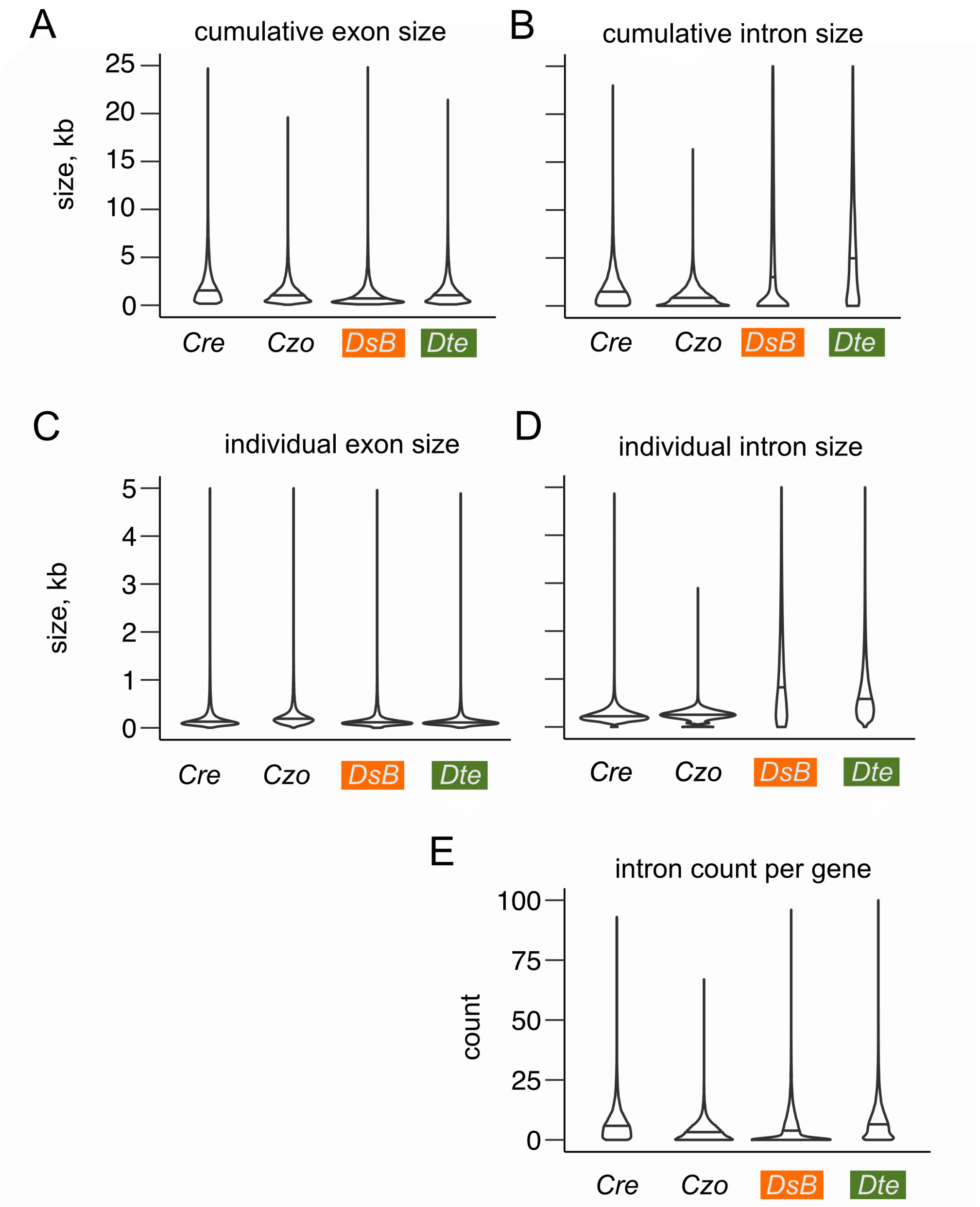


**Fig. S3. Comparison of gene structural features identifies large introns in *Dunaliella*.** The sizes and numbers of exons and introns for all nuclear gene models from four chlorophycean algae were calculated and presented as violin plots as follows: (**A**) the combined length of all protein coding, exonic nucleotides per gene, i.e. the CDS length, (**B**) the combined length of intronic nucleotides per gene, (**C**) individual exon sizes, (**D**) individual intron sizes, and (**E**) the number of introns per gene. Species are indicated in the figure by a three-letter code as follows: *Cre* = *Chl. reinhardtii* (17,741 genes), *Czo* = *Chr. zofingiensis* (15,366 genes), *DsB* = *D. salina* Bardawil (25,074 genes), *Dte* = *D. tertiolecta* (14,351 genes). A horizontal line in each violin represents the median.

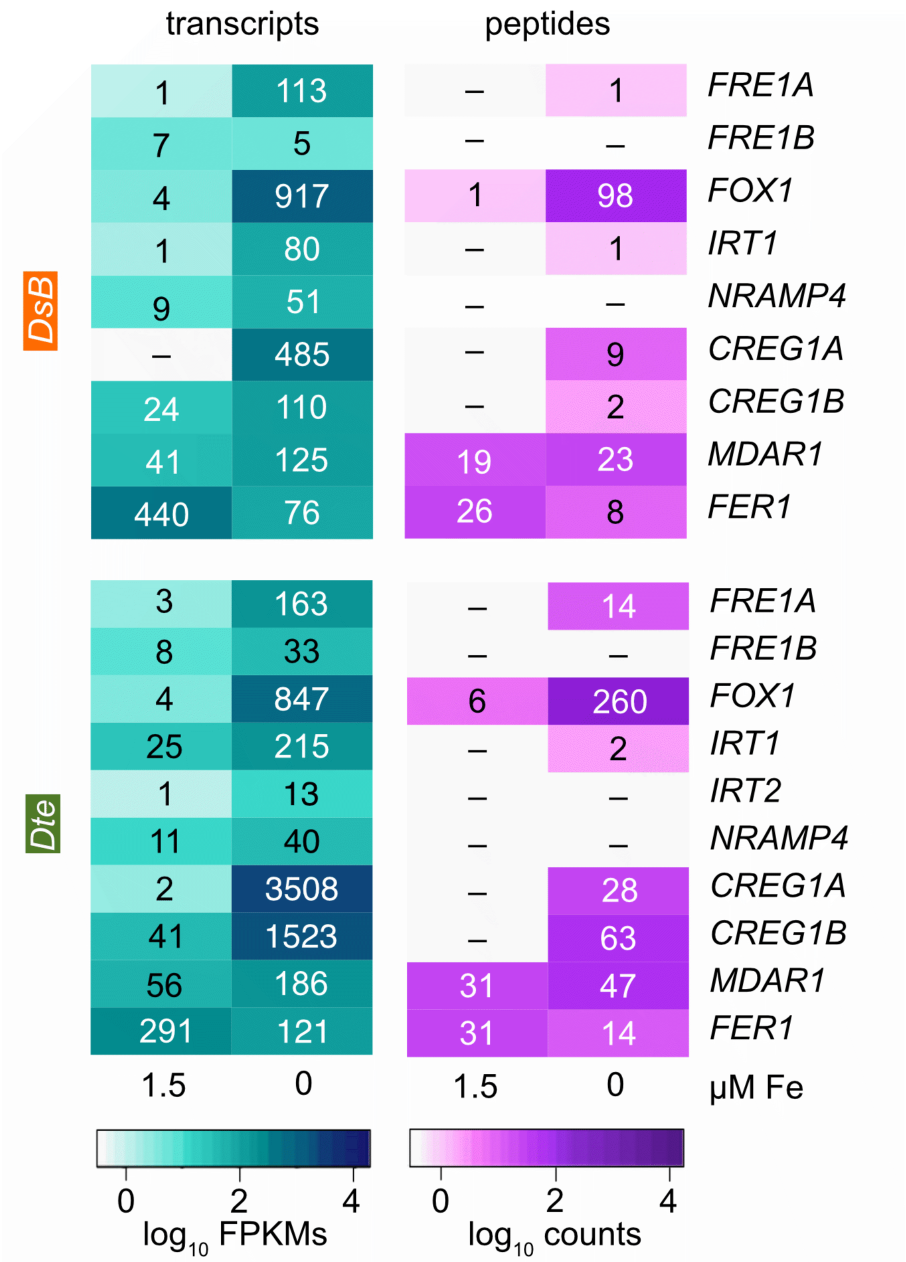


**Fig. S4. Heatmaps of transcript and protein abundance for key Fe-homeostasis genes.** Transcript abundance estimates as $log_{10}$-transformed FPKMs and protein abundance estimates as $log_{10}$-transformed spectral counts for important Fe homeostasis genes are presented as heatmaps for *D. salina* Bardawil and *D. tertiolecta* cultures grown +/– 1.5 μM Fe.

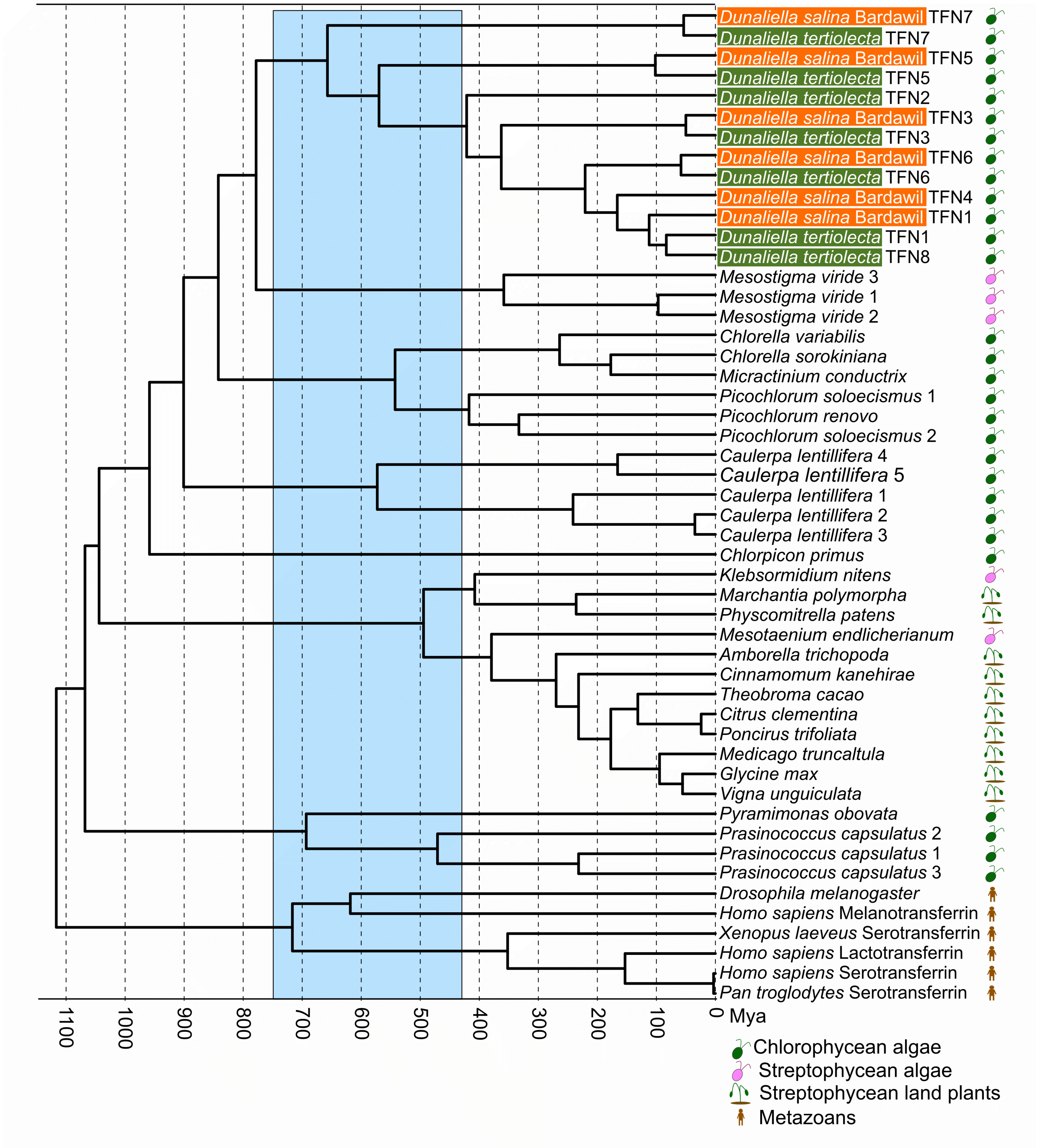


**Fig. S5. Expansion of transferrin proteins in *Dunaliella* and other species**. A chronogram demonstrates the expansion of transferrin proteins in the *Dunaliella* lineage and Viridiplantae more broadly. Select Metazoan transferrin proteins were included as an outgroup. A light blue box indicates the approximate time of the Neoproterozoic oxygenation event.

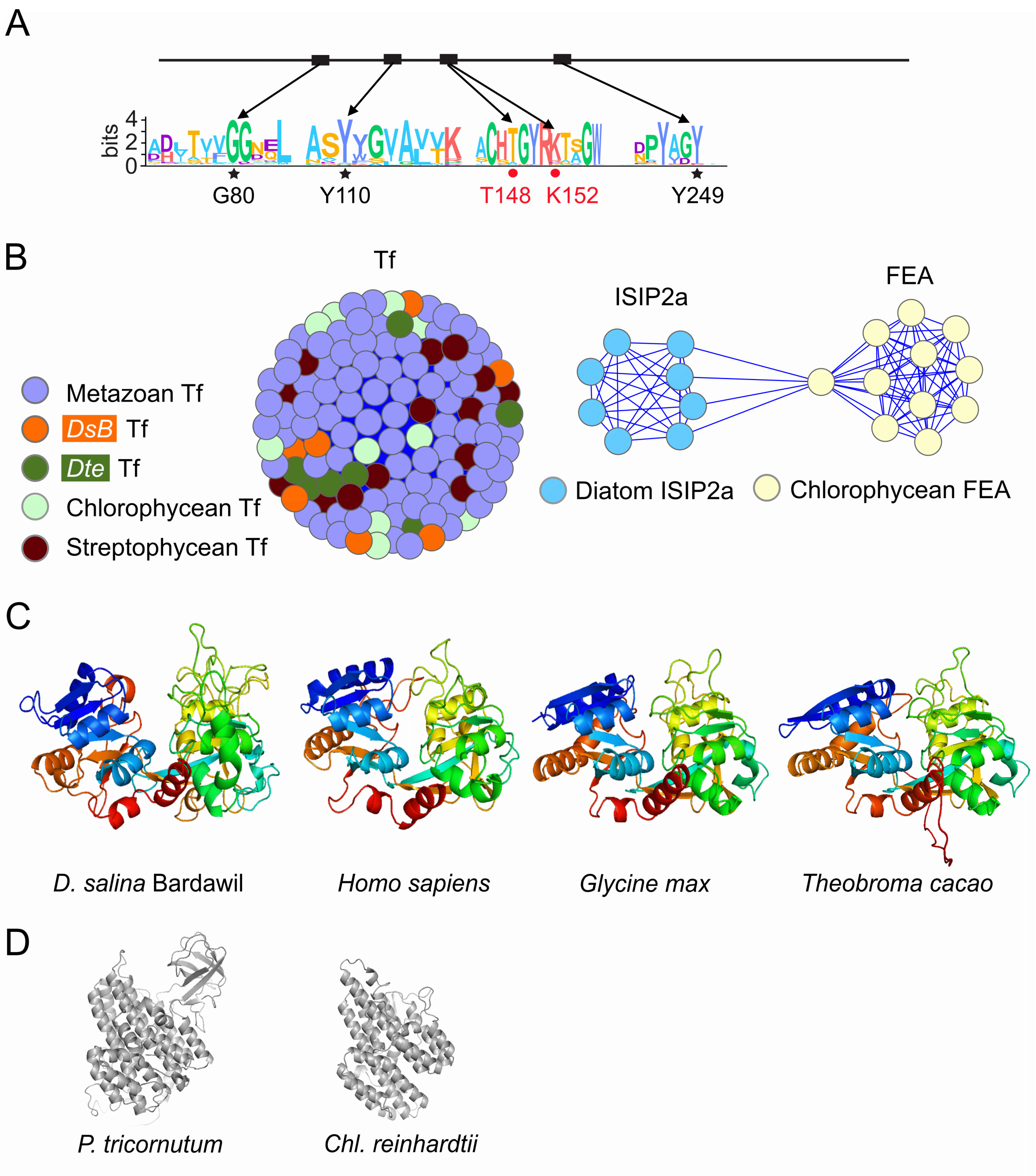


**Fig. S6. Fe-binding proteins: a comparison of transferrins with ISIP2a/FEAs.** (**A**) Regions of high amino acid conservation among the 44 *Dunaliella* Tf domains were identified and are shown here as a logo plot. Highly conserved residues important for Fe and anion binding are indicated as follows: black stars = Fe binding and red circles = anion binding. (**B**) Transferrin (Tf), Fe-assimilation protein (FEA), and Iron induced starvation protein 2a (ISIP2a) from a range of species were subjected to a blastp all-vs-all search using a e-value cutoff of $1 \times 10^{-7}$, and used to construct a protein similarity network. (**C**) Structural predictions of a Tf domain from four species. From left to right: *D. salina* Bardawil TFN1 domain 1, *H. sapiens* STF C-terminal domain, *G. max* XP_006606915 and *T. cacao* XP_007009809. (**D**) Structural predictions of *P. tricornutum* ISIP2a and *Chl. reinhardtii* FEA1.

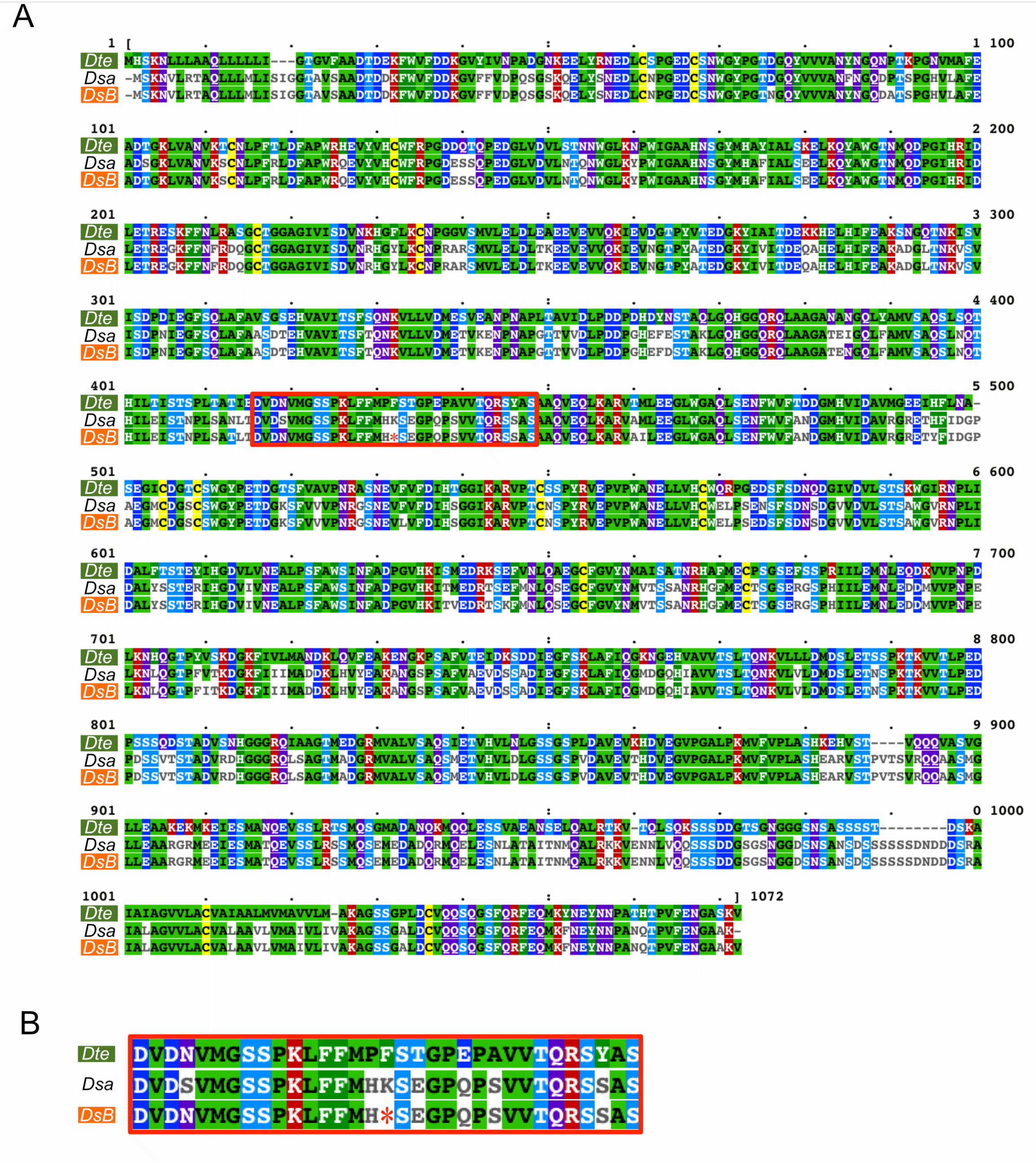


**Fig. S7. Conservation of SUPT1 suggests a recent nonsense mutation in *D. salina* Bardawil.** SUPT1 is a putative siderophore-Fe binding protein in *Dunaliella*. (**A**) Multiple sequence alignment of SUPT1 protein sequences from *D. tertiolecta* (*Dte*), *D. salina* CCAP 19/18 (*Dsa*), and *D. salina* Bardawil LB 2538 (*DsB*) is presented. A red asterisk indicates a single stop codon in *D. salina* Bardawil SUPT1 at position 431 that breaks up the otherwise conserved open reading frame. blastp alignment of *D. salina* Bardawil versus *D. tertiolecta* SUPT1 gives a bit score of 1708 with 822/1052 (78%) identities and 16/1052 (1%) gaps. blastp alignment of *D. salina* Bardawil LB 2538 versus *D. salina* CCAP 19/18 SUPT1 gives a bit score of 2182 with 1054/1070 (99%) identities and 1/1070 (0%) gaps. (**B**) An expanded view of the red-boxed region surrounding the stop codon in *D. salina* Bardawil SUPT1.

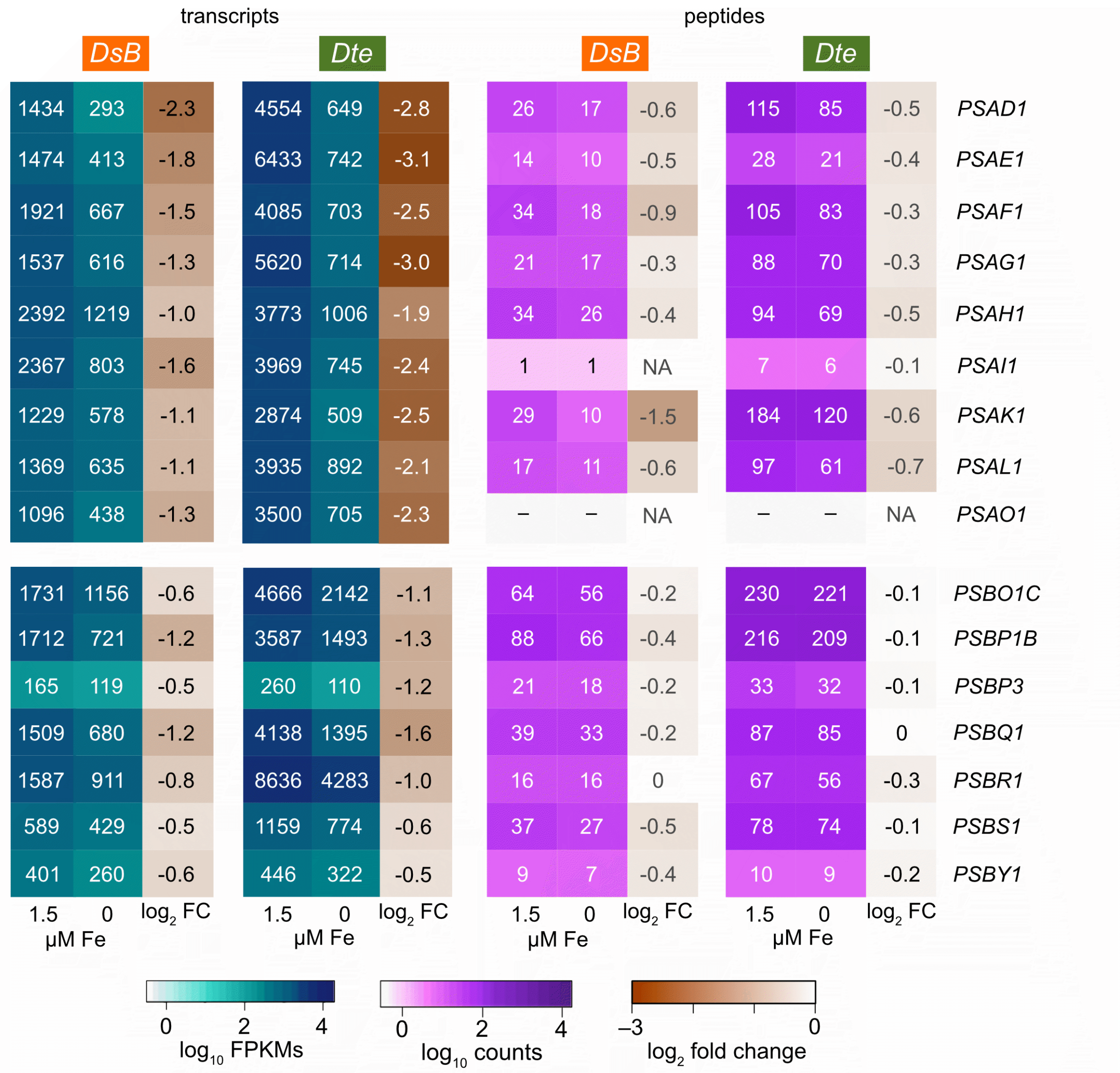


**Fig. S8. Photosystem transcript and polypeptide abundance changes in response to –Fe.** Transcript abundance estimates as $log_{10}$-transformed FPKMs and protein abundance estimates as $log_{10}$-transformed spectral counts for major components of photosystem I (*PSAx*) and photosystem II (*PSBx*) are presented as heatmaps for *D. salina* Bardawil and *D. tertiolecta* cultures grown +/– 1.5 µM Fe. Additionally, the $log_2$ transformed fold change for each comparison is plotted in the third column.

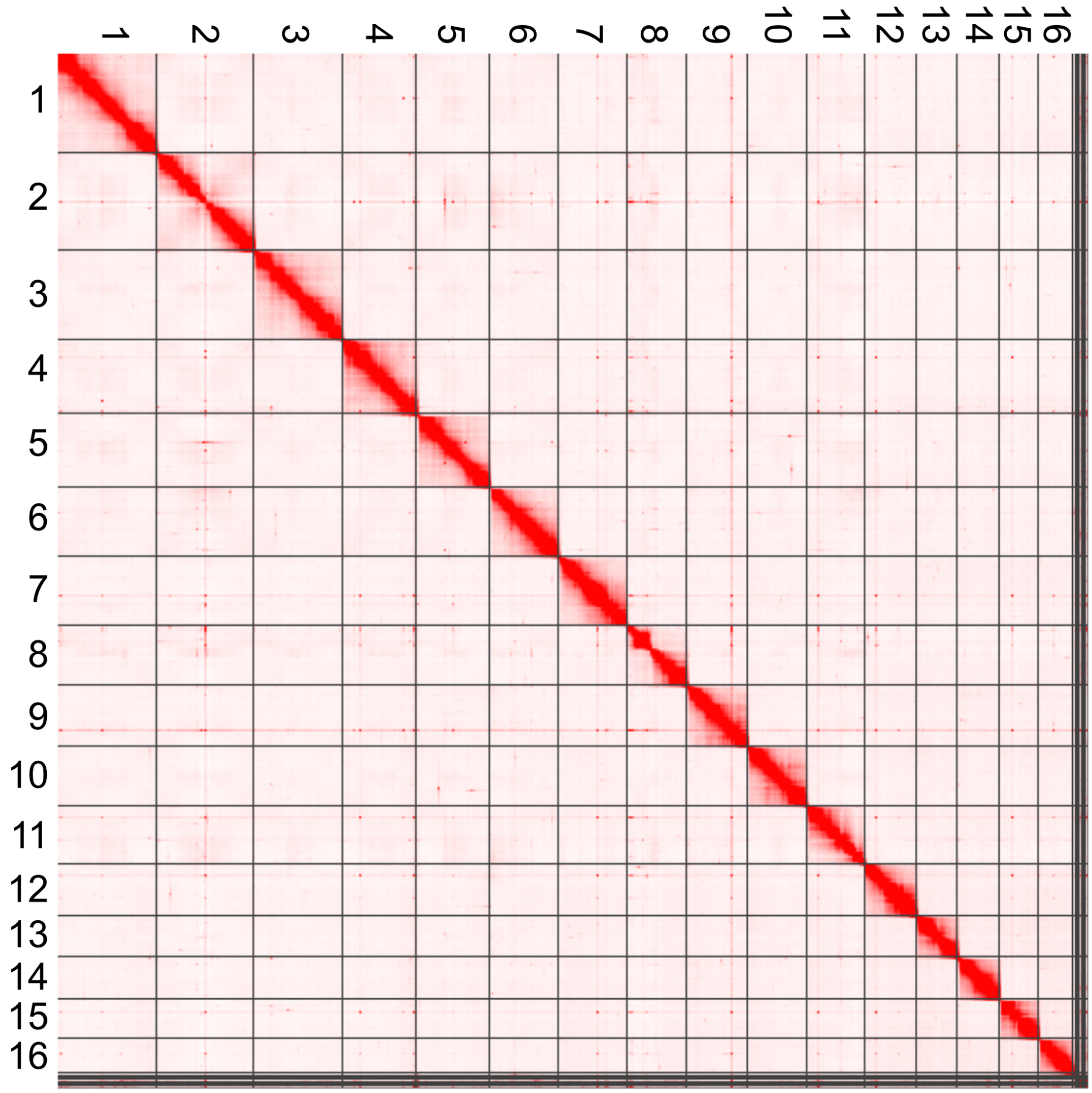


**Fig. S9. The assembled contigs from *D. salina* Bardawil were scaffolded to chromosome scale with the aid of Hi-C sequencing.** Presented here is an interaction map of the Hi-C sequencing data demonstrating organization of the genome into 16 chromosomes.

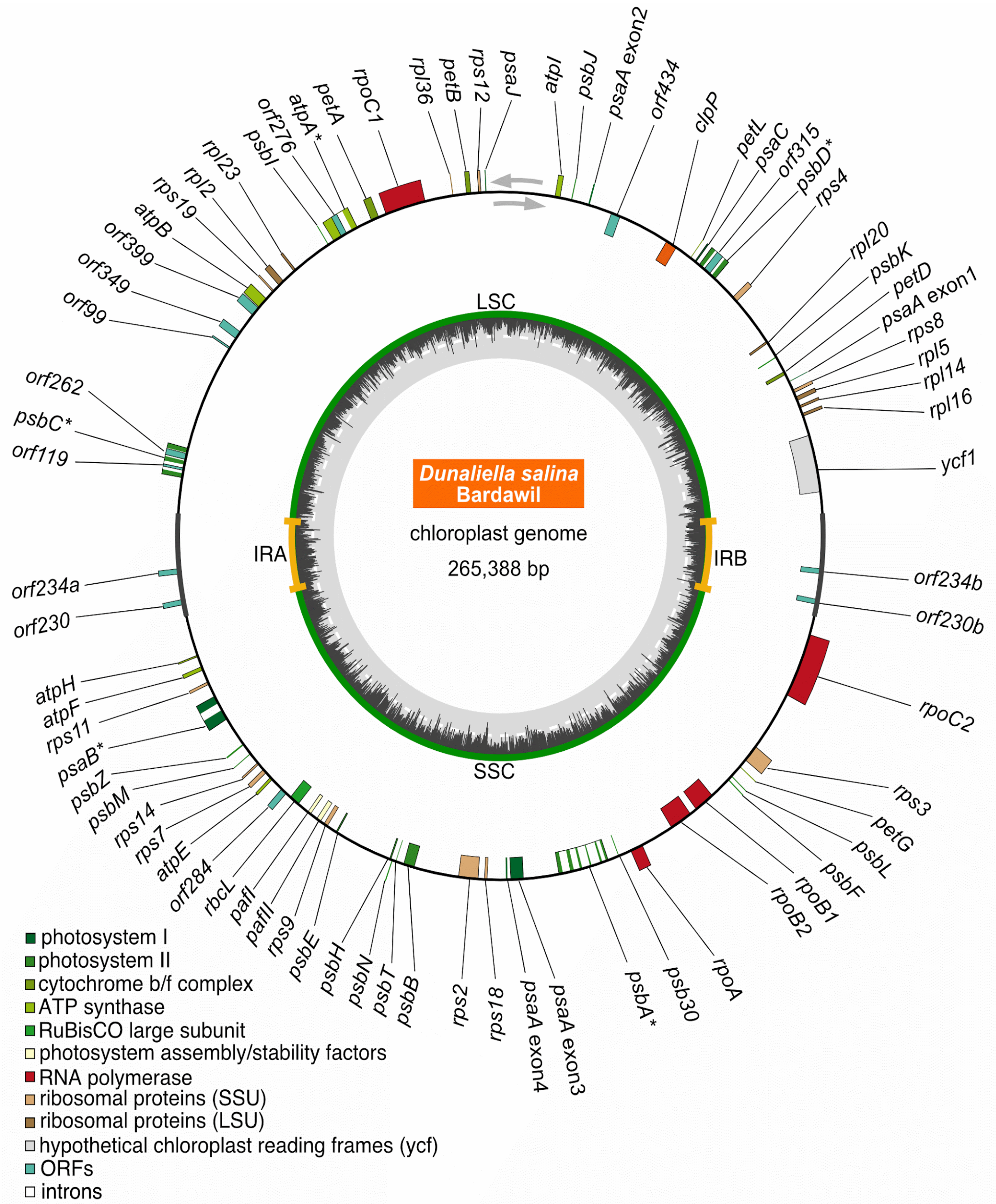


**Figure S10. The circular plastome of *D. salina* Bardawil.** In the inner circle, the GC content is plotted as a histogram in dark gray with a dashed white line indicating 50%. Outside of that, the long single copy (LSC) and short single copy (SSC) regions are shown in green separated by the two inverted repeat regions (IRA and IRB) shown in orange. In the outer circle, the positions of protein coding genes are shown as boxes that are color-coded based on function. Genes expressed in a clockwise fashion are plotted inside the circle and genes expressed counterclockwise are on the outside. An asterisk indicates genes with introns.

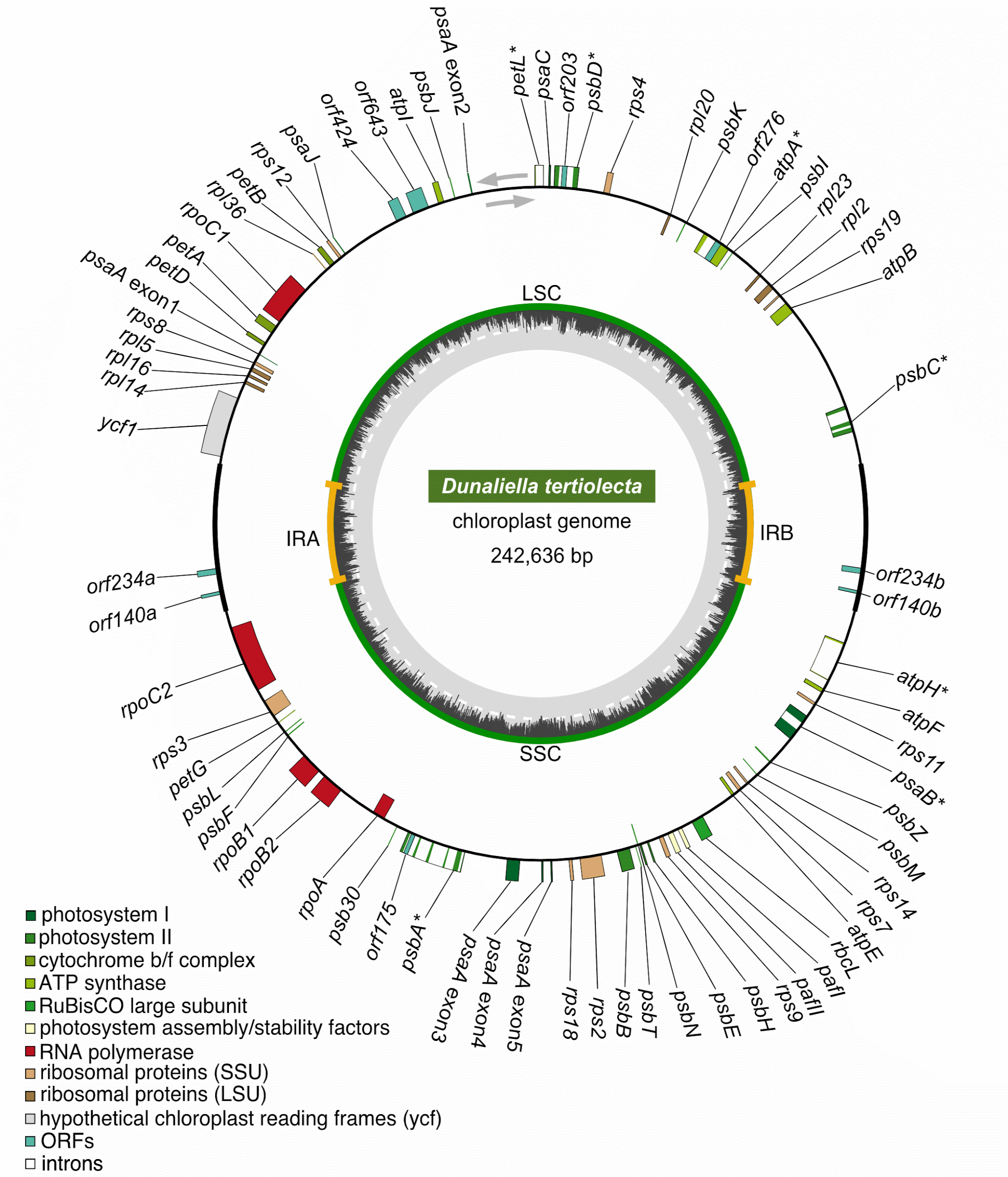


**Figure S11. The circular plastome of *D. tertiolecta*.** In the inner circle, the GC content is plotted as a histogram in dark gray with a dashed white line indicating 50%. Outside of that, the long single copy (LSC) and short single copy (SSC) regions are shown in green separated by the two inverted repeat regions (IRA and IRB) shown in orange. In the outer circle, the positions of protein coding genes are shown as boxes that are color-coded based on function. Genes expressed in a clockwise fashion are plotted inside the circle and genes expressed counterclockwise are on the outside. An asterisk indicates genes with introns.

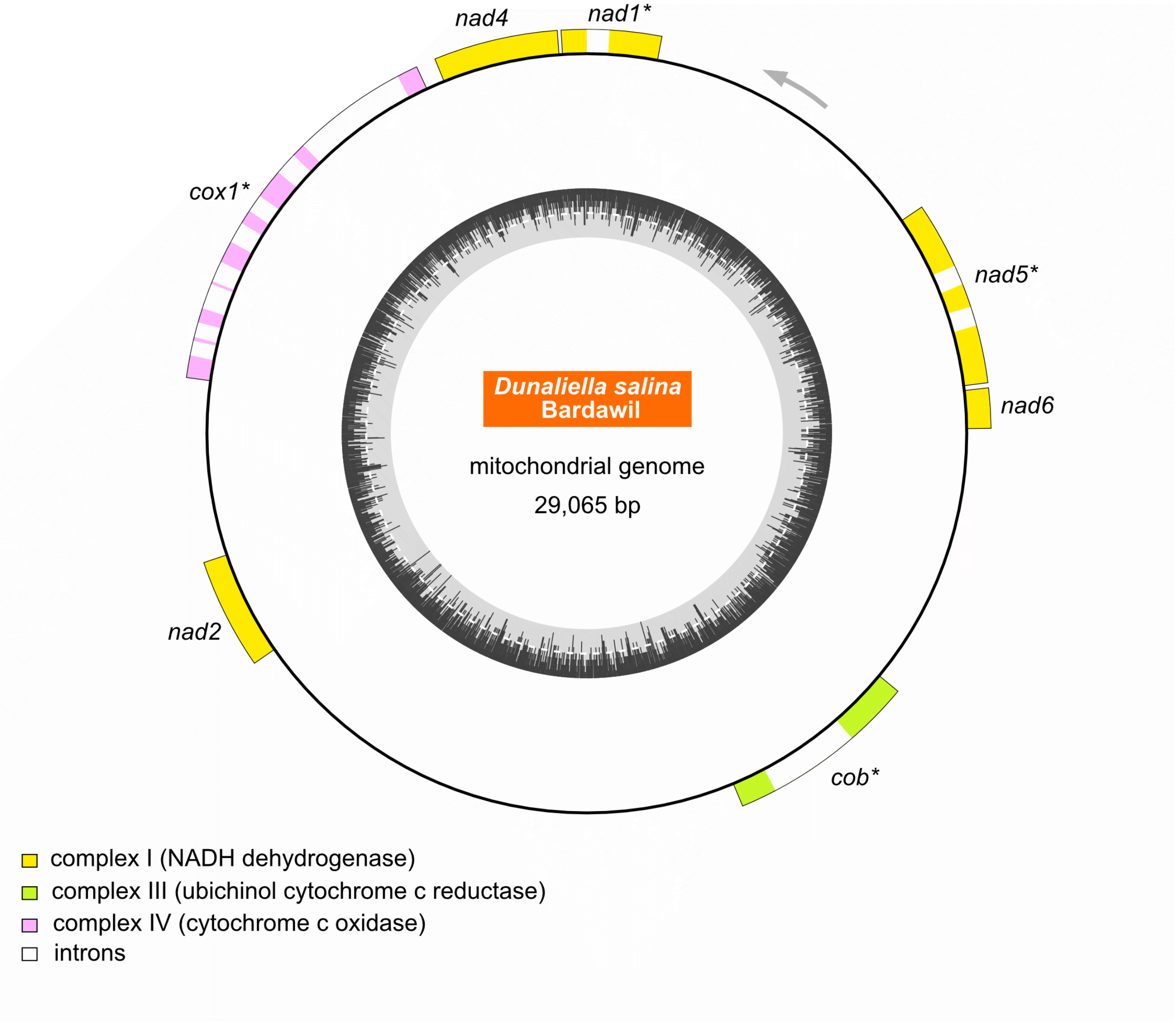


**Figure S12. The circular mitogenome of *D. salina* Bardawil.** In the inner circle, the GC content is plotted as a histogram in dark gray with a dashed white line indicating 50%. In the outer circle, the positions of protein coding genes are shown as boxes that are color-coded based on function. All genes in this map are expressed in a counterclockwise fashion (gray arrow). An asterisk indicates genes with introns.

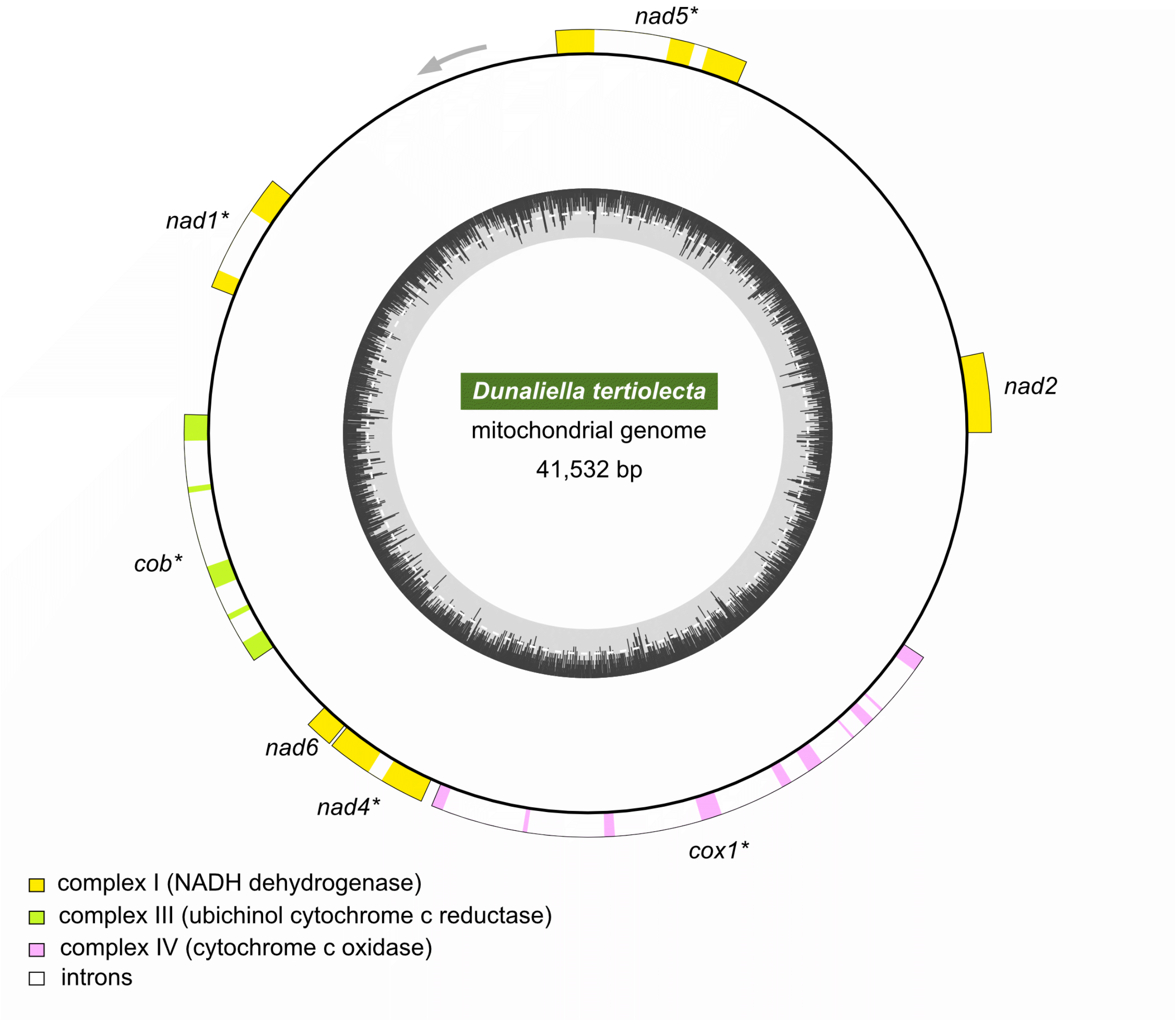


**Figure S13. The circular mitogenome of *D. tertiolecta*.** In the inner circle, the GC content is plotted as a histogram in dark gray with a dashed white line indicating 50%. In the outer circle, the positions of protein coding genes are shown as boxes that are color-coded based on function. All genes in this map are expressed in a counterclockwise fashion (gray arrow). An asterisk indicates genes with introns.

**Supporting tables**

**Table S1. Summary of *Dunaliella* genome repeat contents**

| | | *Dunaliella tertiolecta* | | *Dunaliella salina* Bardawil | |
|---|---|---|---|---|---|
| **Repeat** | | **Length (Mb)** | **Genome (%)** | **Length (Mb)** | **Genome (%)** |
| **LINE** | All | 44.18 | 23.29 | 63.97 | 17.63 |
| | *Dualen* | 27.73 | 14.62 | 46.30 | 12.76 |
| | *RTE-X* | 16.57 | 8.73 | 16.04 | 4.42 |
| | *L1* | 0.10 | 0.05 | 0.52 | 0.14 |
| **SINE** | | 1.05 | 0.55 | 1.32 | 0.36 |
| **LTR** | All | 3.25 | 1.71 | 18.46 | 5.09 |
| | *Gypsy* | 1.41 | 0.74 | 14.61 | 4.03 |
| | *Copia* | 0.25 | 0.13 | 0.66 | 0.18 |
| **DIRS** | *PAT-like* | 0.13 | 0.07 | 0.23 | 0.06 |
| **DNA transposon** | All | 6.86 | 3.62 | 35.99 | 9.92 |
| | *EnSpm* | 1.37 | 0.72 | 9.41 | 2.59 |
| | *hAT* | 0.68 | 0.36 | 1.81 | 0.50 |
| | *Harbinger* | 0.18 | 0.10 | 0.39 | 0.11 |
| **Crypton** | *CryptonF* | 0.00 | 0.00 | 3.64 | 1.00 |
| **Unknown** | | 13.80 | 7.27 | 36.59 | 10.09 |
| **Tandem repeats** | All | 17.52 | 9.23 | 59.09 | 16.29 |
| | microsatellites | 3.86 | 2.04 | 25.75 | 7.10 |
| | satellites | 13.66 | 7.20 | 33.34 | 9.19 |
| **Total** | | **86.36** | **45.53** | **218.56** | **60.25** |

**Table S2. Conservation of Residues in Transferrin Domains**

| | | | Fe binding | | | anion binding | |
|---|---|---|---|---|---|---|---|
| | | | **G80*** | **Y110†** | **Y249‡** | **T148§** | **K152¶** |
| ***D. salina* Bardawil** | TFN1 | 1 | - | - | - | - | - |
| | | 2 | - | - | - | - | - |
| | | 3 | - | - | - | - | - |
| | | 4 | - | - | - | - | - |
| | TFN3 | 1 | **A** | **A** | - | - | **R** |
| | | 2 | - | - | - | - | - |
| | TFN4 | 1 | - | - | - | - | - |
| | TFN5 | 1 | - | - | - | - | - |
| | TFN6 | 1 | - | - | - | - | - |
| | | 2 | - | - | - | - | - |
| | | 3 | - | - | - | - | - |
| | | 4 | - | - | - | - | - |
| | | 5 | - | - | - | - | - |
| | | 6 | - | - | - | - | - |
| | | 7 | - | - | - | - | - |
| | | 8 | - | - | - | - | - |
| | | 9 | - | - | - | - | - |
| | TFN7 | 1 | - | **G** | - | - | **G** |
| | | 2 | - | - | **E** | - | **Q** |
| | | 3 | - | - | - | - | - |
| ***D. tertiolecta*** | TFN1 | 1 | - | - | - | - | - |
| | | 2 | - | - | - | - | - |
| | | 3 | - | - | - | - | - |
| | TFN2 | 1 | - | - | - | - | **R** |
| | | 2 | **I** | - | **R** | **M** | **R** |
| | TFN3 | 1 | **A** | **A** | - | - | **R** |
| | | 2 | - | - | - | - | - |
| | TFN5 | 1 | - | - | - | - | - |
| | TFN6 | 1 | - | - | - | - | - |
| | | 2 | - | - | - | - | - |
| | | 3 | - | - | - | - | - |
| | | 4 | - | - | - | - | - |
| | | 5 | - | - | - | - | - |
| | | 6 | - | - | - | - | - |
| | | 7 | - | - | - | - | - |
| | | 8 | - | - | - | - | - |
| | | 9 | - | - | - | - | - |
| | | 10 | - | - | - | - | - |
| | TFN7 | 1 | - | - | - | - | - |
| | | 2 | - | - | - | - | - |
| | | 3 | - | - | - | - | - |
| | TFN8 | 1 | - | **H** | - | **A** | **T** |
| | | 2 | - | - | - | **A** | - |
| | | 3 | - | - | - | **A** | **E** |

Listed residues are based on the amino acid position in *D. tertiolecta* TFN1 domain 1.
Dashes represent conserved amino acids, variants are indicated.

* Equivalent to D63 in *H. sapiens* STF N-lobe (12)

† Equivalent to Y95 in *H. sapiens* STF N-lobe

‡ Equivalent to Y188 in *H. sapiens* STF N-lobe

§ Equivalent to T120 in *H. sapiens* STF N-lobe

¶ Equivalent to R124 in *H. sapiens* STF N-lobe

### Table S3. Accession Information for Other Species Analyzed

| Species | Strain / Isolate / Ecotype | Short Label* | Version | Data Type(s) | Source† | Ref. |
|---|---|---|---|---|---|---|
| ***Dunaliella salina*** | CCAP 19/18 | *Dsa* | v1.0 | genome<br>transcriptome<br>proteome | Phytozome | (3) |
| ***Dunaliella tertiolecta*** | LB 999 | *Dte*-C | N/A | transcriptome<br>proteome | SI materials | (2) |
| ***Dunaliella tertiolecta*** | LB 999 | *Dte*-L | N/A | transcriptome<br>proteome | SI materials | (1) |
| ***Dunaliella salina*** | TG | N/A | N/A | transcriptome<br>proteome | GEO GSE97378 | (13) |
| ***Dunaliella salina*** | HG01 | N/A | N/A | transcriptome<br>proteome | GEO GSE120965 | (14) |
| ***Dunaliella viridis*** | CCAP19/3 | N/A | N/A | proteome | GEO GSE74466 | (15) |
| ***Chlamydomonas priscuii*** | UWO 241 | N/A | v1.0 | proteome | Phycocosm | (9) |
| ***Chlamydomonas reinhardtii*** | CC-503 | *Cre* | v5.6 | genome<br>proteome | Phytozome | (16) |
| | | | | expression | GEO GSE35305 | (17) |
| ***Volvox carteri*** | HK10 | *Vca* | v2.1 | genome<br>proteome | Phytozome | (18) |
| ***Chromochloris zofingiensis*** | SAG 211-14 | *Czo* | v5.2.3.2 | genome<br>proteome | Phytozome | (19) |
| ***Auxenochlorella protothecoides*** | UTEX 25 | *Apr* | N/A | proteome | SRA PRJNA289168 | unpub. |
| ***Coccomyxa subellipsoidea*** | C-169 | *Csu* | v2.0 | proteome | Phytozome | (20) |
| ***Gonium pectorale*** | NIES-2863 | *Gpe* | v1.0 | genome<br>proteome | Phycocosm | (21) |
| ***Tetrabaena socialis*** | NIES-571 | *Tso* | v1.0 | genome<br>proteome | Phycocosm | (22) |
| ***Arabidopsis thaliana*** | Col-0 | *Ath* | TAIR10 | genome<br>proteome | Phytozome | (23) |
| | | | | expression | GEO GSE163190 | (24) |
| ***Monoraphidium neglectum*** | SAG 48-87 | *Mne* | v1.0 | genome<br>proteome | Phycocosm | (25) |
| ***Chlamydomonas eustigmata*** | NIES-2499 | *Ceu* | v1.0 | genome<br>proteome | Phycocosm | (26) |
| ***Raphidocelis subcapitata*** | NIES-35 | *Rsu* | v1.0 | genome<br>proteome | Phycocosm | (27) |
| ***Saccharomyces cerevisiae*** | S288C | *Sce* | 64-3-1 | genome<br>proteome | SGD | (28) |
| ***Oryza sativa*** | Nipponbare/<br>japonica | *Osa* | v7.0 | proteome | Phytozome | (29) |
| | | | | expression | GEO GSE131238 | (30) |
| ***Fragilariopsis cylindrus*** | CCMP 1102 | *Fcy* | v1.0 | proteome | Phycocosm | (31) |
| | | | | expression | ArrayExpress E-MTAB-5024 | (32) |
| ***Phaeodactylum tricornutum*** | CCAP 1055 | *Ptr* | v2.0 | proteome | Phycocosm | (33) |
| | | | | expression | GEO GSE8675 | (34) |
| ***Thalassiosira pseudonana*** | CCMP 1335 | *Tps* | v3.0 | genome<br>proteome | Phycocosm | (35) |
| ***Thalassiosira oceanica*** | CCMP 1005 | *Toc* | v1.0 | proteome | Phycocosm | (36) |
| | | | | expression | SRA PRJNA382002 | (37) |

* Short label as used in figures, if applicable.
† Data source:
Phytozome = https://phytozome-next.jgi.doe.gov/
Phycocosm = https://phycocosm.jgi.doe.gov/
GEO = https://www.ncbi.nlm.nih.gov/geo/
SRA = https://www.ncbi.nlm.nih.gov/sra/
ArrayExpress = https://www.ebi.ac.uk/biostudies/arrayexpress/studies
SGD = https://www.yeastgenome.org/

**Table S4. BUSCO Scores**

| | Species | single | duplicated | fragmented | missing |
|---|---|---|---|---|---|
| Genome | *Dunaliella tertiolecta* LB 999* | 1442<br>94.9% | 21<br>1.4% | 9<br>0.6% | 47<br>3.1% |
| | *Dunaliella salina* Bardawil LB 2538* | 1421<br>93.5% | 22<br>1.4% | 14<br>0.9% | 62<br>4.1% |
| | *Dunaliella salina* CCAP 19/18 (3) | 1284<br>84.5% | 15<br>1.0 | 52<br>3.4% | 168<br>11.1% |
| Transcriptome | *Dunaliella tertiolecta* LB 999* | 1368<br>90.1% | 18<br>1.2% | 13<br>0.9% | 120<br>4.1% |
| | *Dunaliella tertiolecta* LB 999 *de novo* transcriptome Cho group (2) | 929<br>61.2% | 201<br>13.2% | 65<br>4.3% | 324<br>21.3% |
| | *Dunaliella tertiolecta* LB 999 *de novo* transcriptome Lee group (1) | 859<br>56.6% | 317<br>20.9% | 99<br>6.5% | 244<br>16.1% |
| | *Dunaliella salina* Bardawil LB 2538* | 1309<br>86.2% | 113<br>7.4% | 19<br>1.3% | 78<br>5.1% |
| | *Dunaliella salina* CCAP 19/18 (3) | 1128<br>74.3% | 78<br>5.1% | 93<br>6.1% | 220<br>14.5% |

* This work

**Table S5. Assembly Details**

| Species | Size, bp | GC, % | Ns | Total contigs | Total scaf. | N50 (bp)<br>L50 (#)<br>contigs | N50 (bp)<br>L50 (#)<br>scaffolds |
|---|---|---|---|---|---|---|---|
| ***Dunaliella salina*** **Bardawil LB 2538 (v1.1)*** | 363,064,759 | 49.4 | 384,490<br>(0.11%) | 1,003 | 211 | 1,016,134<br>106 | 23,405,360<br>7 |
| ***Dunaliella tertiolecta*** **LB 999 (v1.1)*** | 189,986,397 | 49.4 | 181,322<br>(0.096%) | 96 | 20 | 4,336,382<br>14 | 17,075,954<br>6 |
| ***Dunaliella salina*** **CCAP 19/18 (v1.0)** (3) | 343,704,438 | 49.1 | 62,867,987<br>(18.3%) | 58,282 | 5,509 | 7,229<br>11,486 | 353,034<br>310 |
| ***Chlamydomonas reinhardtii*** **CC-4532 (v6.1)** (38) | 114,631,715 | 64.0 | 1,114,981<br>(0.97%) | 127 | 60 | 2,649,501<br>15 | 6,954,842<br>7 |

* This work

**Table S6. Gene Annotation Details**

| Species | total genes | nuclear genes | plastid genes | mitochondrial genes |
|---|---|---|---|---|
| ***Dunaliella salina*** **Bardawil LB 2538 (v1.1)*** | 25,074 | 24,988 | 79 | 7 |
| ***Dunaliella tertiolecta*** **LB 999 (v1.1)*** | 14,351 | 14,268 | 76 | 7 |
| ***Dunaliella salina*** **CCAP 19/18 (v1.0)** (3) | 16,697 | 16,697 | not included | not included |
| ***Chlamydomonas reinhardtii*** **CC-4532 (v6.1)** (38) | 17,823 | 17,741 | 74 | 8 |

* This work

**Legends for supporting datasets**

**Dataset S1 (separate file). Dunaliella_expression_data.xlsx.** All gene annotations, transcriptomic expression data, and proteomic expression data for *D. salina* Bardawil and *D. tertiolecta*.
